# Cortical wiring by synapse-specific control of local protein synthesis

**DOI:** 10.1101/2021.11.12.468364

**Authors:** Clémence Bernard, David Exposito-Alonso, Martijn Selten, Stella Sanalidou, Alicia Hanusz-Godoy, Fazal Oozeer, Patricia Maeso, Beatriz Rico, Oscar Marín

## Abstract

Neurons use local protein synthesis as a mechanism to support their morphological complexity, which requires independent control across multiple subcellular compartments including individual synapses. However, to what extent local translation is differentially regulated at the level of specific synaptic connections remains largely unknown. Here, we identify a signaling pathway that regulates the local synthesis of proteins required for the formation of excitatory synapses on parvalbumin-expressing (PV^+^) interneurons in the mouse cerebral cortex. This process involves the regulation of the mTORC1 inhibitor Tsc2 by the receptor tyrosine kinase ErbB4, which enables the local control of mRNA translation in a cell type-specific and synapse-specific manner. Ribosome-associated mRNA profiling reveals a molecular program of synaptic proteins that regulates the formation of excitatory inputs on PV^+^ interneurons downstream of ErbB4 signaling. Our work demonstrates that local protein translation is regulated at the level of specific connections to control synapse formation in the nervous system.

---

The extraordinary diversity of animal behaviors relies on the precise assembly of neuronal circuits, a process in which synapse formation plays a critical role. In the cerebral cortex, dozens of different types of excitatory glutamatergic pyramidal cells and inhibitory γ-aminobutyric acid-containing (GABAergic) interneurons are wired through highly specific connectivity motifs [1, 2]. For example, layer 4 excitatory neurons receive inputs from excitatory thalamic neurons and project to layer 2/3 excitatory neurons and feed-forward interneurons [3]. Synapse specificity is established during development by dedicated transcriptional programs [4–10], but whether regulation of mRNA translation is also involved in this process remains to be elucidated.

Protein synthesis is controlled by several pathways [11, 12], including the mechanistic target of rapamycin complex 1 (mTORC1), a molecular complex composed of mTOR kinase and several other proteins that is activated by nutrients and growth factor signals, and inhibited by the proteins Tsc1 and Tsc2 [13, 14]. Multiple mTORC1 pathway proteins, as well as the translation machinery, have been identified in developing axons [15–17], and local protein synthesis occurs at both excitatory and inhibitory synapses in the adult brain [18–24]. To what extent local translation is differentially regulated in closely related cell types and at the level of specific synapses during the wiring of cortical circuits is not known.

## Specific synaptic defects in interneurons lacking Tsc2

To explore the role of protein synthesis in the wiring of different cell types in the cerebral cortex, we generated mice in which we deleted *Tsc2* from the two largest groups of cortical GABAergic interneurons, parvalbumin (PV) and somatostatin (SST) expressing cells. To this end, we crossed *Lhx6-Cre* mice, which drives recombination in early postmitotic PV^+^ and SST^+^ interneurons, with mice carrying conditional (i.e., Cre-dependent) *Tsc2* alleles (*Tsc2^F/F^*) and a reporter for the visualization of recombined cells (see Methods). We chose these two cell types because although they derive from common progenitors in the medial ganglionic eminence (MGE) and the preoptic area (POA), populate the same layers of the neocortex, and are both reciprocally connected with pyramidal cells [25], they end up playing very different roles in cortical information processing [26]. We first confirmed that loss of *Tsc2* leads to overactivation of mTOR signaling in PV^+^ and SST^+^ interneurons by analyzing the levels of phosphorylated S6 ribosomal protein (P-S6rp), a critical downstream effector of mTORC1 [12]. We observed increased levels of P-S6rp in conditional *Tsc2* mutants compared to controls (Suppl. Fig. 1A,B). We also found that the cell size of PV^+^ and SST^+^ interneurons is significantly larger in conditional *Tsc2* mutants than controls (Suppl. Fig. 1A-C), reinforcing the notion that mTOR signaling is indeed overactive in these cells [27, 28].

We next investigated whether loss of Tsc2 affects synapse formation onto PV^+^ and SST^+^ interneurons in the neocortex. To this end, we assessed the number of excitatory synapses received by cortical PV^+^ and SST^+^ interneurons by quantifying puncta containing vesicular glutamate transporter 1 (VGluT1), a characteristic component of excitatory glutamatergic terminals, and PSD95, the major scaffolding protein in the excitatory postsynaptic density. We found that loss of one and, even more so, two *Tsc2* alleles in PV^+^ interneurons led to a prominent increase in the density of excitatory synapses received by these cells compared to control littermates (Fig. 1A,B). In contrast, conditional deletion of *Tsc2* caused no changes in the density of excitatory synapses received by SST+ interneurons (Fig. 1C,D). This differential impact on PV^+^ and SST^+^ synaptic wiring was confirmed by recording spontaneous excitatory postsynaptic currents (sEPSCs) in both cell types (Fig. 1E). *Tsc2* deletion in PV^+^ interneurons led to a significant increase in the frequency of sEPSCs with no changes in their amplitude (Fig. 1F,G), while neither the frequency nor the amplitude of sEPSCs were changed in SST^+^ interneurons lacking *Tsc2* (Fig. 1H,I). We also observed cell type-specific changes in the intrinsic properties of PV^+^ and SST^+^ interneurons following the conditional deletion of *Tsc2* (Fig. 2A). These changes led to reduced excitability in PV^+^ interneurons in conditional *Tsc2* mutants compared to controls, while no difference was observed for SST^+^ interneurons (Suppl. Fig. 2B). Altogether, these results revealed that loss of *Tsc2* differentially impacts PV^+^ and SST^+^ interneurons, suggesting that the role of Tsc2 in synapse formation is cell type specific.

**Fig. 1.**
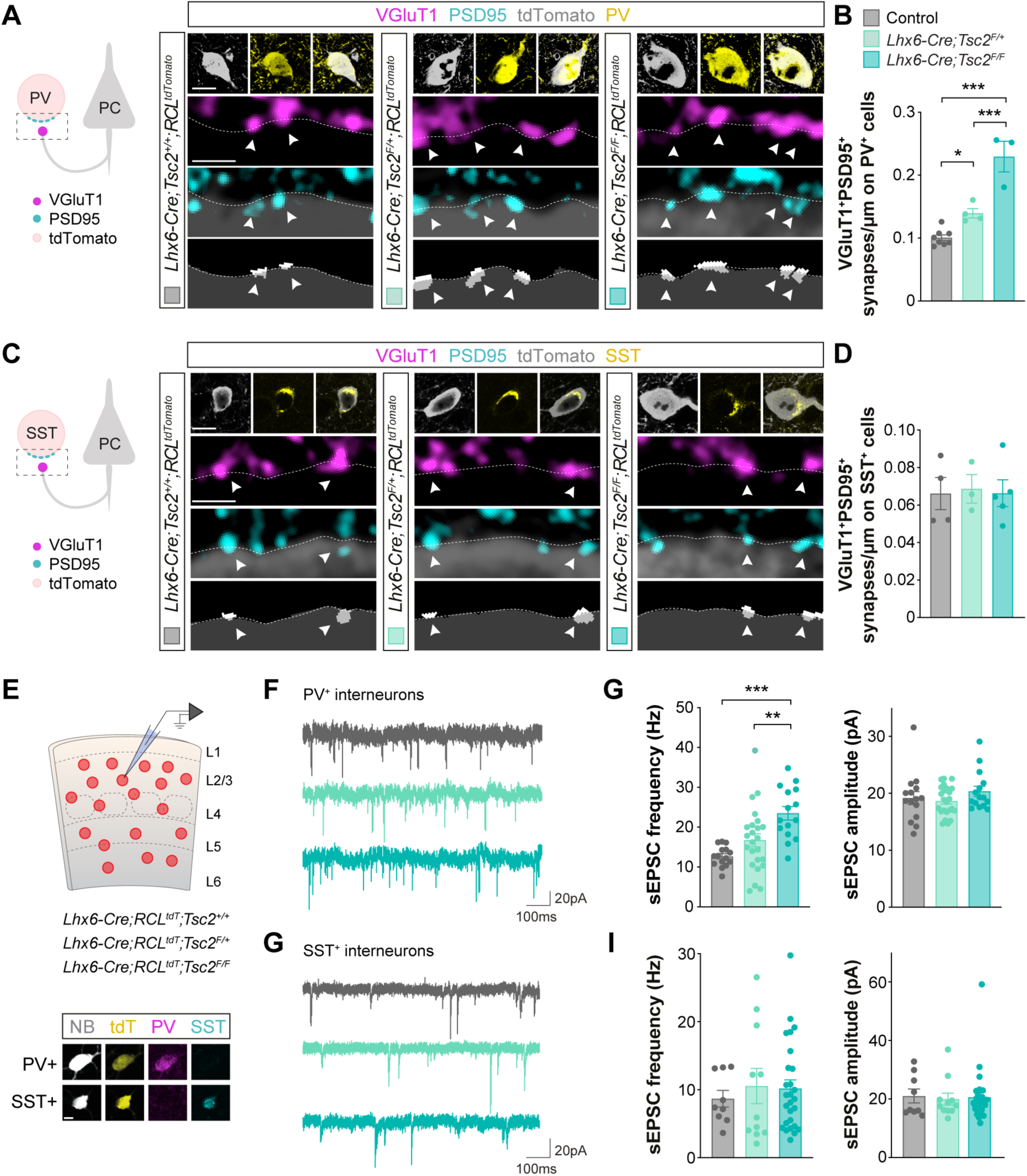
Conditional deletion of *Tsc2* from PV^+^ and SST^+^ interneurons differentially impact excitatory inputs received by these cells. (**A**) Schematic of synaptic markers analyzed (left). Confocal images (top) and binary images (bottom) illustrating presynaptic VGluT1^+^ puncta (magenta) and postsynaptic PSD95^+^ clusters (cyan) in PV^+^ (yellow) tdTomato^+^ (grey) interneurons from P18-21 control, heterozygous and homozygous conditional *Tsc2* mutants. (**B**) Quantification of the density of VGluT1^+^PSD95^+^ synapses contacting PV^+^ interneurons (control, *n* =111 cells from 8 mice; heterozygous*, n* = 70 cells from 4 mice; homozygous*, n* = 46 cells from 3 mice). (**C**) Schematic of synaptic markers analyzed (left). Confocal images (top) and binary images (bottom) illustrating presynaptic VGluT1^+^ puncta (magenta) and postsynaptic PSD95^+^ clusters (cyan) in SST^+^ (yellow) tdTomato^+^ (grey) interneurons from P18-21 control, heterozygous and homozygous conditional *Tsc2* mutants. (**D**) Quantification of the density of VGluT1^+^PSD95^+^ synapses contacting SST^+^ interneurons (control, *n* = 67 cells from 4 mice; heterozygous*, n* = 41 cells from 3 mice; homozygous*, n* = 60 cells from 5 mice). (**E**) Schematic of experimental design (top) and post-recording labelling of neurobiotin (NB, grey) -filled tdTomato^+^ (yellow) cells with PV (magenta) and SST (cyan) (bottom). (**F**) Example traces of sEPSCs recorded from PV^+^ interneurons from P18-21 control, heterozygous and homozygous conditional *Tsc2* mutants. (**G**) Quantification of the frequency (left) and amplitude (right) of sEPSCs from PV^+^ interneurons (control *n* =15 cells from 5 mice, *Lhx6-Cre;Tsc2^F/+^* n=24 cells from 8 mice, *Lhx6-Cre;Tsc2^F/F^ n* =15 cells from 6 mice). (**H**) Example traces of sEPSCs recorded from SST^+^ interneurons from P18-21 control, heterozygous and homozygous conditional *Tsc2* mutants. (**I**) Quantification of the frequency (left) and amplitude (right) of sEPSCs from SST^+^ interneurons (control, *n* = 9 cells from 5 mice; heterozygous*, n* = 11 cells from 9 mice; homozygous*, n* = 27 cells from 7 mice). One-way ANOVA followed by Tukey’s multiple comparisons test: *P < 0.05, **P < 0.01, ***P < 0.001. Data are mean ± s.e.m. Scale bar, 10 *µ*m and 1 *µ*m (high magnification) (A, C), and 10 *µ*m (E).

**Fig. 2.**
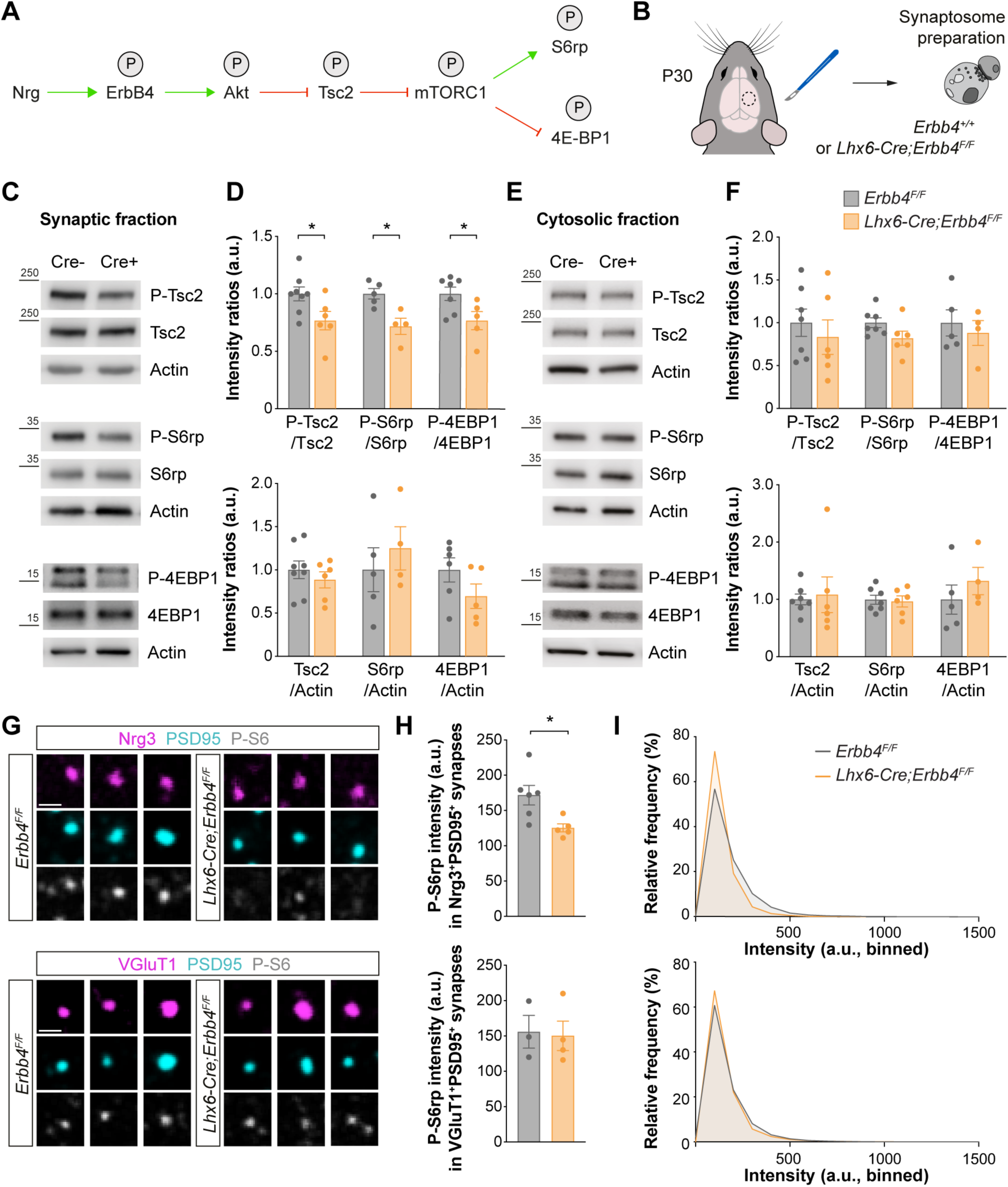
ErbB4 regulates mTOR signaling at excitatory synapses received by PV^+^ interneurons. (**A**) Hypothetical signaling pathway. P indicates phosphorylation, green arrow activation, red arrow inhibition. (**B**) Schematic of experimental design. (**C**) Phosphorylation and protein expression of Tsc2, S6rp, 4EBP1 and actin assessed by Western blot of cortical synaptic fractions from P30 homozygous conditional *Erbb4* mice and their control littermates. (**D**) Quantification of phosphorylation of Tsc2, S6rp and 4EBP1 normalized to the total expression of the corresponding protein (top). Quantification of expression levels of Tsc2, S6rp and 4EBP1 normalized to actin (bottom) (Tsc2: control*, n* = 8 mice, homozygous*, n* = 6 mice; S6rp: control*, n* = 5 mice, homozygous*, n* = 4 mice; 4EBP1: control*, n* = 7 mice, homozygous*, n* = 5 mice). (**E**) Phosphorylation and protein expression of Tsc2, S6rp, 4EBP1 and actin assessed by Western blot of cortical cytosolic fractions from P30 homozygous conditional *Erbb4* mice and their control littermates. (**F**) Quantification of phosphorylation of Tsc2, S6rp and 4EBP1 normalized to the total expression of the corresponding protein (top). Quantification of expression levels of Tsc2, S6rp and 4EBP1 normalized to actin (bottom) (Tsc2 and S6rp: control*, n* = 7 mice, homozygous*, n* = 6 mice; 4EBP1: control*, n* = 5 mice, homozygous*, n* = 4 mice). (**G**) Confocal images illustrating phosphorylation of S6rp (P-S6rp, grey) in Nrg3^+^ (magenta) PSD95^+^ (cyan) synaptosomes (top) and in VGluT1^+^ (magenta) PSD95^+^ (cyan) synaptosomes (bottom) from P21 homozygous conditional *Erbb4* mice and their control littermates. (**H**) Quantification of P-S6rp staining intensity in Nrg3^+^PSD95^+^ (top) and VGluT1^+^PSD95^+^ synaptosomes (bottom). (**I**) Relative frequency distribution of P-S6rp staining intensity in Nrg3^+^PSD95^+^ synaptosomes (top) and in VGluT1^+^PSD95^+^ synaptosomes (bottom) (Nrg3^+^PSD95^+^: control*, n* = 22,063 synaptosomes from 6 mice, homozygous*, n* = 18,050 synaptosomes from 5 mice; VGluT1^+^PSD95^+^: control*, n* = 30,774 synaptosomes from 3 mice, homozygous*, n* = 31,547 synaptosomes from 4 mice). Two-tailed Student’s unpaired t-tests: *P < 0.05. Data are mean ± s.e.m. Scale bar, 1 *µ*m.

We next wondered whether the function of Tsc2 in synapse development might be synapse-type specific. We reasoned that if Tsc2 plays a general role in synapse formation in PV^+^ interneurons, loss of Tsc2 should also affect the development of the synapses made by these cells onto pyramidal cells. We used synaptotagmin-2 (Syt2) to specifically identify the presynaptic compartment of PV^+^ basket cell synapses [29] and gephyrin (Geph), a postsynaptic scaffolding protein of GABAergic synapses. Unexpectedly, we found no differences in the density of Syt2^+^Geph^+^ synaptic puncta contacting the soma of pyramidal cells in conditional *Tsc2* mutants compared to control littermates (Suppl. Fig. 3A,B). Thus, although the loss of Tsc2 causes a global disruption of mTOR signaling in PV^+^ and SST^+^ cells (e.g., increased cell size and P-S6rp), Tsc2 seems to be uniquely involved in the wiring of interneurons in a cell-type and synapse-type specific manner.

## Tsc2 functions downstream of ErbB4 signaling in synapses

Since the loss of Tsc2 function only leads to changes in the excitatory synaptic input of PV^+^ interneurons, we reasoned that Tsc2 activity could be regulated locally by a signaling pathway specific to these synapses and necessary for their formation. In other cellular contexts, Tsc2 activity is inhibited by signaling pathways involved in stimulating cell growth through the phosphorylation of Akt [14, 30]. The receptor tyrosine kinase ErbB4 is specifically required for the formation of excitatory synapses onto PV^+^ interneurons [31–33], can activate Akt via PI3K phosphorylation [34], and interacts with Tsc2 in synaptosome preparations obtained from the mouse neocortex during synaptogenesis (Suppl. Fig. 4). This led us to hypothesize that ErbB4 functions upstream of Tsc2 to regulate its activity during the development of these specific synapses (Fig. 2A). To begin testing this hypothesis, we analyzed Tsc2 phosphorylation in synaptosomes obtained from the neocortex of control and interneuron specific *Erbb4* conditional mutants (Fig. 2B). We found reduced Akt-mediated phosphorylation of Tsc2 in cortical synapses from *Erbb4* conditional mutants compared to controls (Fig. 2C,D). We also found reduced phosphorylation of the mTORC1 effectors S6rp and eIF4E-binding protein 1 (4E-BP1) in synaptosomes from *Erbb4* conditional mutants (Fig. 2C,D), which indicates that the loss of ErbB4 function decreases mTORC1 synaptic activity due to the overactivation of Tsc2. Remarkably, none of these changes were observed in cytosolic fractions (Fig. 2E,F), which suggests that ErbB4 is specifically required to modulate Tsc2 signaling at the synapse.

We next investigated whether the mTORC1 inactivation observed in *Erbb4* conditional mutants occurs at specific cortical synapses. Since ErbB4 is only expressed by specific classes of interneurons in the cerebral cortex [31, 35], we hypothesized that changes would be limited to synapses that would normally contain ErbB4 receptors, such as the excitatory synapses received by PV^+^ basket cells. To test this idea, we isolated and plated cortical synaptosomes and analyzed the phosphorylation of S6rp in postsynaptic structures identified with PSD95. We identified excitatory synapses onto inhibitory neurons using neuregulin 3 (Nrg3), which is specifically present in presynaptic excitatory terminals contacting PV^+^ interneurons [36, 37]. We found decreased P-S6rp specifically in Nrg3^+^/PSD95^+^ synaptosomes from *Erbb4* conditional mutants compared to controls, but no differences in ErbB4-independent glutamatergic synaptosomes (Fig. 2G,I). These results indicate that ErbB4 uniquely regulates mTOR signaling in excitatory synapses received by PV^+^ interneurons.

To add further support to the idea that ErbB4 and Tsc2 function in the same signaling pathway controlling the formation of excitatory synapses onto PV^+^ cells, we performed a genetic interaction experiment. We generated *Erbb4* conditional mutants carrying a conditional *Tsc2* allele to compromise the function of Tsc2 in PV^+^ interneurons lacking ErbB4 and measured the density of excitatory synapses received by these cells. We found that deleting one *Tsc2* allele from PV^+^ cells is sufficient to rescue the loss of excitatory synapses found in *Erbb4* conditional mutants (Suppl. Fig. 5A,B). Altogether, these results demonstrate that Tsc2 functions downstream of ErbB4 in regulating the excitatory synaptic input of PV^+^ interneurons.

## Loss of ErbB4 leads to specific changes in synaptic translatome

To identify the specific targets of ErbB4 that are involved in the development of synapses in PV^+^ interneurons, we first obtained the synaptic translatome of MGE-derived interneurons from control and *Erbb4* conditional mutants. To this end, we bred into these mice alleles carrying a mutation in the locus encoding the ribosomal protein L22 (Rpl22) that allows the conditional tagging of ribosomes with a hemagglutinin epitope (HA) tag [38]. We then prepared synaptosomes from the neocortex of P15 mice, pulled-down ribosomes from this preparation using anti-HA beads to isolate interneuron-specific ribosome-associated mRNA transcripts in control and *ErbB4* conditional mutants, and analyzed them by RNA sequencing (Fig. 3A and Suppl. Fig. 6A,D).

**Fig. 3.**
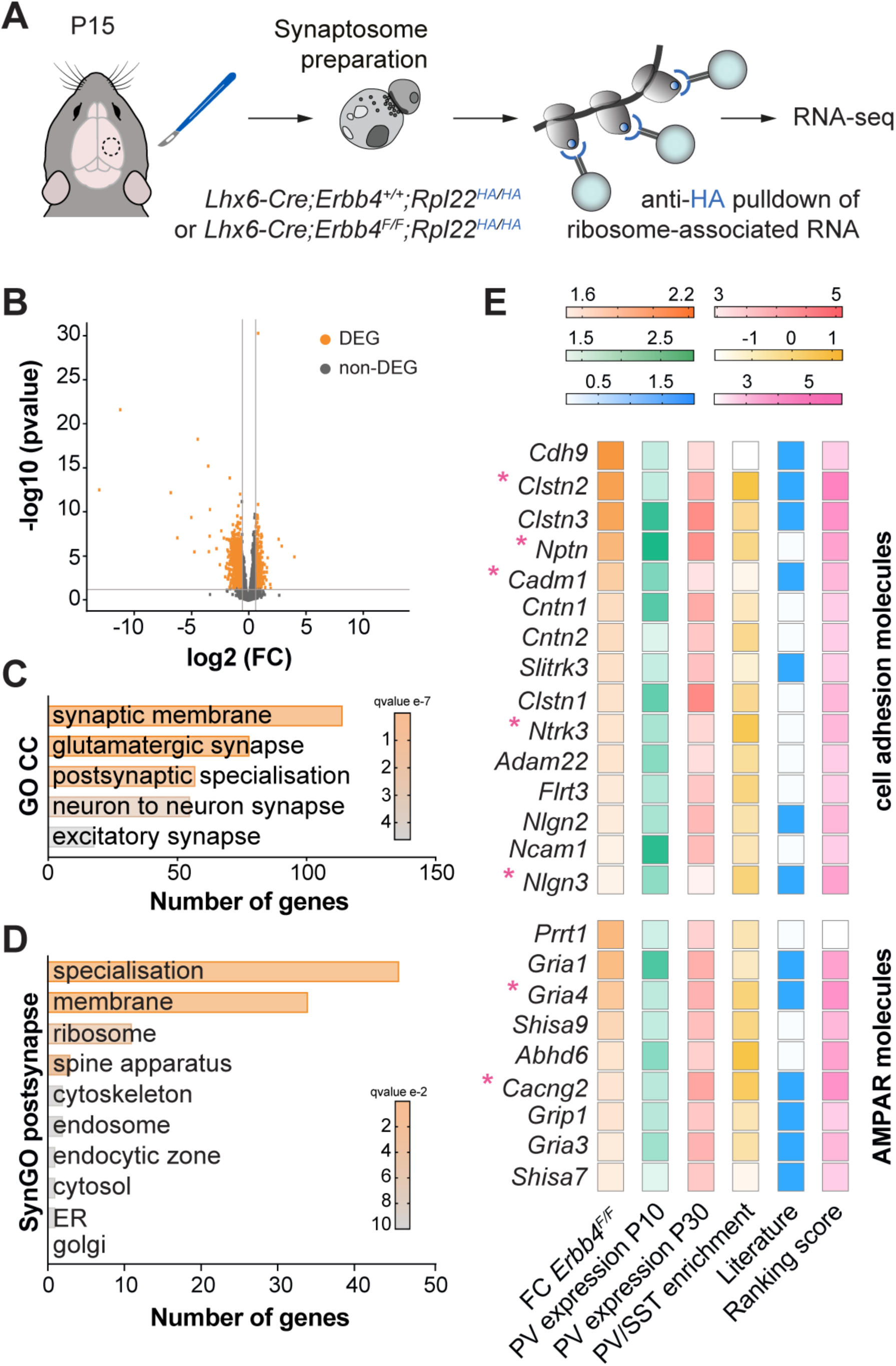
Synaptic ribosome-associated mRNA transcripts are altered in *Erbb4* conditional mutants. (**A**) Schematic of experimental design. (**B**) Volcano plot displaying significantly differentially expressed ribosome associated RNAs (DEG, orange) in P15 cortical synaptosomes from homozygous conditional *Erbb4* mice compared to controls. Each dot represents one gene. FC: fold-change (FC > 1.5, P < 0.05). (**C**) Selected Gene Ontology (GO) Cellular Components (CC) terms significantly enriched in the dataset of downregulated genes in homozygous conditional *Erbb4* mutants compared to controls. (**D**) Synaptic Gene Ontology (SynGO) postsynaptic cellular component categories significantly enriched in the dataset of downregulated genes in homozygous conditional *Erbb4* mutants compared to controls. (**E**) Heatmaps showing the selection criteria for 15 “cell adhesion molecules” genes and 9 “AMPA receptors” genes. The asterisks indicate genes selected for validation.

We found that 70% of the differentially expressed genes were downregulated in *Erbb4* conditional mutants compared to controls (Fig. 3B and Suppl. Fig. 6D). Gene Ontology (GO) analysis revealed an enrichment in genes involved in processes such as “synapse organization”, “neurotransmitter receptor activity” and “glutamatergic synapse” (Fig. 3C and Suppl. Fig. 6E), highlighting synaptic alterations in *Erbb4* conditional mutants. We then identified genes coding for proteins with postsynaptic localization and/or function using Synaptic Gene Ontology (SynGO) annotations [39]. This analysis revealed that, within the postsynaptic category, the most enriched terms were “postsynaptic specialization” and “postsynaptic membrane”, and more specifically genes encoding cell adhesion molecules and AMPA receptors (Fig. 3C,D and Suppl. Fig. 6F). Using a set of four additional criteria (see Methods), including their relative enrichment in PV^+^ cells, we selected seven candidates for functional validation: five cell adhesion molecules, SynCAM1 (encoded by *Cadm1*), Nptn, Nlgn3, TrkC (encoded by *Ntrk3*) and Clstn2, and two AMPA receptor-related proteins, GluA4 (encoded by *GriA4*) and Stargazin (encoded by *Cacng2*) (Fig. 3E). We confirmed that all these mRNAs are expressed by cortical PV^+^ interneurons during the period of synaptogenesis (Suppl. Fig. 7A,B). In addition, we found that these proteins cluster at the surface of PV^+^ cells in close apposition to innervating axon terminals expressing the presynaptic ErbB4 ligand Nrg3 (Suppl. Fig. 7C,E). Altogether, our data reveal dysregulation of postsynaptic molecular complexes in PV^+^ cells lacking ErbB4.

## ErbB4 regulates local protein synthesis at the synapse

Having identified ribosome-associated mRNAs that might be critical for the formation of excitatory synapses onto PV^+^ cells, we next wondered whether ErbB4 may regulate this process by modulating the local translation of these transcripts. mTORC1 signaling is critical for the modulation of protein synthesis [12], and our previous results revealed that mTORC1 synaptic activity decreases in the absence of ErbB4 (Fig. 2). ErbB4 could therefore regulate synapse formation by controlling protein synthesis at the synapse through the inhibition of Tsc2. To test this hypothesis, we assessed whether activation of ErbB4 signaling in cortical synaptosomes increases the translation of the ribosome-associated mRNAs we have previously identified to be downregulated in *ErbB4* conditional mutants. To this end, we treated cortical synaptosomes with a soluble form of the epidermal growth factor (EGF)-like signaling domain of neuregulin to activate ErbB4 receptors [40] and examined the levels of the candidate proteins by Western blot (Fig. 4A). We found increased protein levels for all but one of the downstream candidates of ErbB4 signaling (Fig. 4B,C). This effect was abolished for 5 out of 7 candidates by the protein synthesis inhibitor cycloheximide (Suppl. Fig. 8A-C), which demonstrated that ErbB4 signaling induces the translation of these proteins at the synapse. Altogether, these experiments revealed that ErbB4 regulates the local translation of synaptic proteins.

**Fig. 4.**
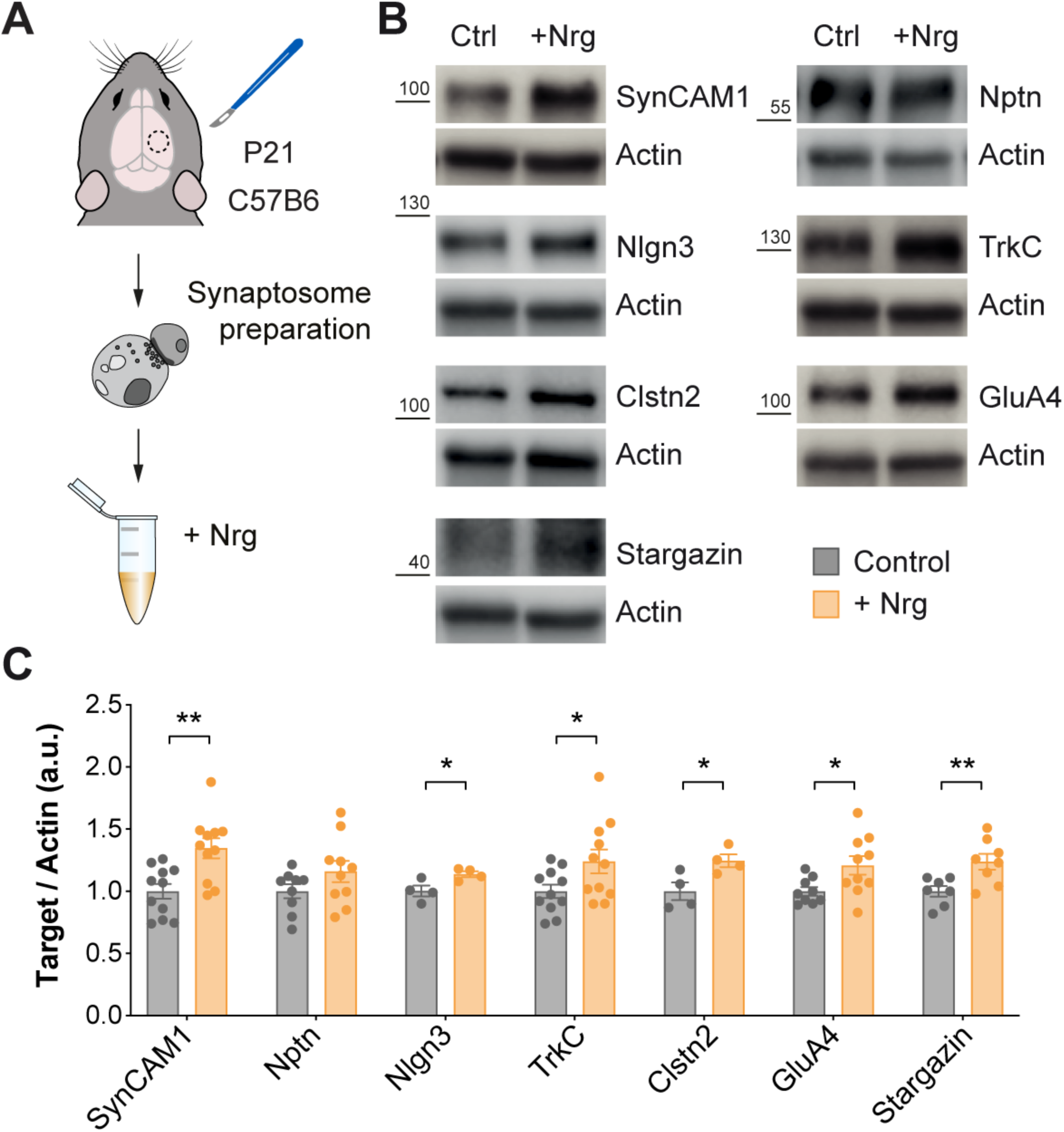
ErbB4 signaling at synapses regulates protein levels of synaptic proteins. (**A**) Schematic of experimental design. (**B**) Protein expression of SynCAM1, Nptn, Nlgn3, TrkC, Clstn2, GluA4, Stargazin and actin assessed by Western blot of cortical synaptic fractions treated with neuregulin (Nrg) from P21 C57B6 mice. (**C**) Quantification of expression levels of SynCAM1, Nptn, Nlgn3. TrkC, Clstn2, GluA4 and Stargazin normalized to actin. One-tailed Student’s unpaired t-tests: *P < 0.05, **P < 0.01 (SynCAM1 and TrkC: control, *n* =11 synaptosomes: +Nrg, *n* = 11 synaptosomes; Nptn and GluA4: control, *n* = 9 synaptosomes, +Nrg, *n* = 10 synaptosomes; Nlgn3 and Clstn2: control, *n* = 4 synaptosomes, +Nrg, *n* = 4 synaptosomes; Stargazin: control, *n* = 7 synaptosomes, +Nrg, *n* = 8 synaptosomes). Data are mean ± s.e.m.

## Molecular mediators of excitatory synapse formation on PV+ interneurons

To assess whether the proteins being translated downstream of ErbB4 are indeed involved in the formation of excitatory synapses on cortical PV^+^ interneurons, we performed interneuron-specific loss-of-function experiments *in vivo* using a conditional gene knockdown strategy [41]. In brief, we designed conditional short-hairpin RNA vectors to target the target genes (*shCadm1, shNptn*, *shNlgn3*, *shNtrk3*, *shClstn2, shGria4*, and *shCacng2*, with *shLacZ* as a control) and confirmed their effectiveness in vitro (Suppl. Fig. 9A). We generated adeno-associated viruses (AAV) expressing the most effective shRNA constructs, injected them into the neocortex of *Lhx6-Cre* neonates, and confirmed their ability to downregulate the expression of the corresponding target genes in vivo (Suppl. Fig. 9B). We then assessed the number of excitatory synapses received by PV^+^ interneurons expressing control and experimental *shRNAs* (Fig. 5A). We found that, compared to controls, reducing the expression of each of the seven targets in PV^+^ interneurons led to a decrease in the density of excitatory synapses received by these cells at P21 (Fig. 5B-D). These results revealed a complex molecular program regulated by ErbB4 that includes SynCAM1, Nptn, Nlgn3, TrkC, Clstn2, GluA4 and Stargazin, and controls the formation of excitatory synapses onto PV^+^ interneurons (Suppl. Fig. 10).

**Fig. 5.**
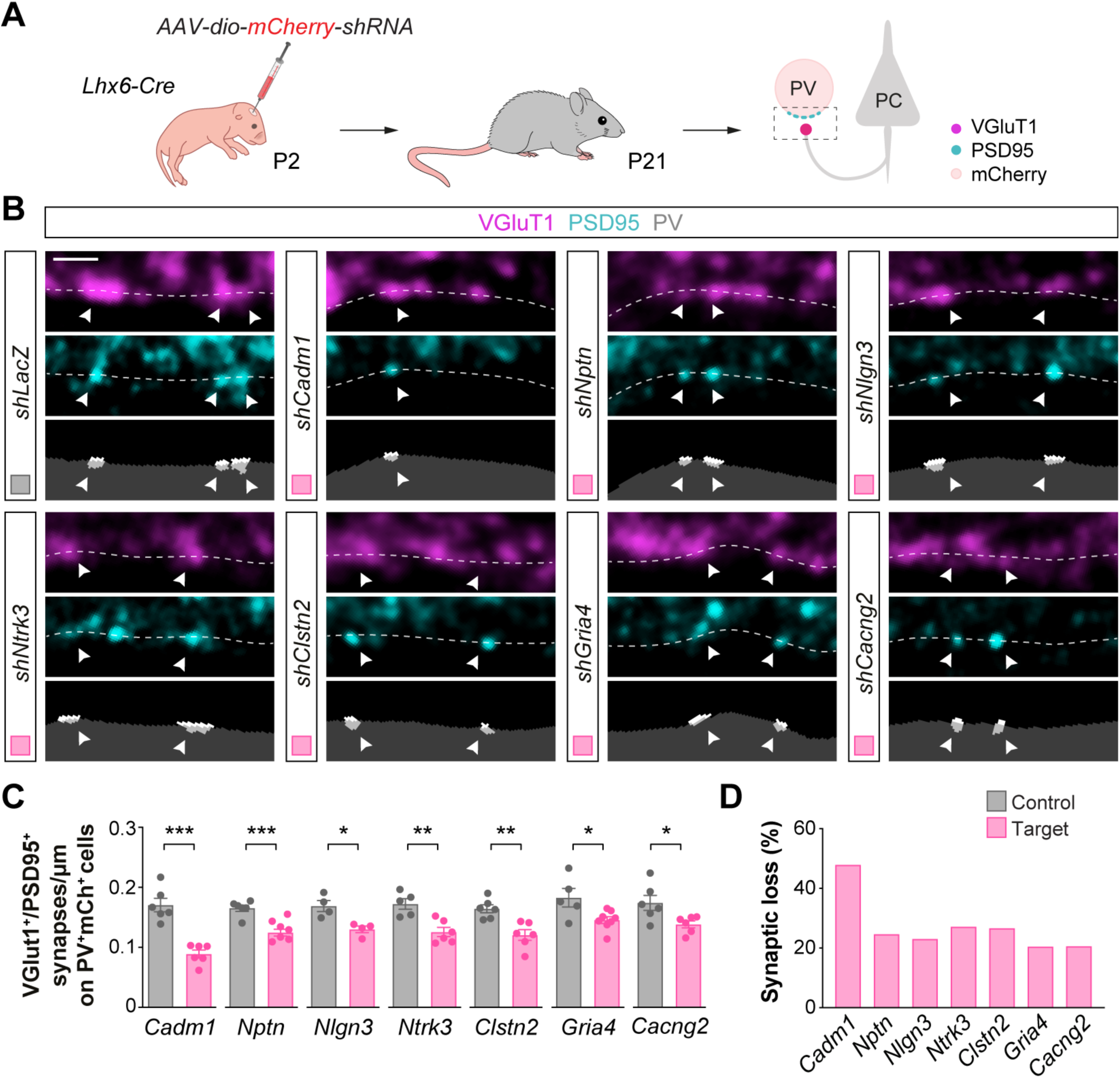
ErbB4 targets control the formation of excitatory inputs onto PV^+^ interneurons. (**A**) Schematic of experimental design. (**B**) Confocal images (top) and binary images (bottom) illustrating presynaptic VGluT1^+^ puncta (magenta) and postsynaptic PSD95^+^ clusters (cyan) in PV^+^ interneurons (grey) from P21 *Lhx6-Cre* mice injected with viruses expressing *shRNAs* targeting the genes of interest or with a control virus (*shLacZ*). (**C**) Quantification of the density of VGluT1^+^PSD95^+^ synapses contacting PV^+^ interneurons in knockdown and control mice. Two-tailed Student’s unpaired t-tests: *P < 0.05, **P < 0.01, ***P < 0.001; *shCadm1* (*n* = 122 cells from 6 mice) and control (*n* = 113 cells from 6 mice); *shNptn* (*n* = 131 cells from 6 mice) and control (*n* = 119 cells from 6 mice); *shNlgn3* (*n* = 86 cells from 4 mice) and control (*n* = 68 cells from 4 mice); *shNtrk3* (*n* = 126 cells from 6 mice) and control (*n* = 105 cells from 5 mice); *shClstn2* (*n* = 120 cells from 6 mice) and control (*n* = 112 cells from 6 mice); *shGria4* (*n* = 121 cells from 8 mice) and control (*n* = 93 cells from 5 mice); *shCacng2* (*n* = 115 cells from 6 mice) and control (*n* = 109 cells from 6 mice). (**D**) Proportion of synaptic loss in PV^+^ interneurons upon knockdown of ErbB4 downstream targets. Data are mean ± s.e.m. Scale bar, 1 *µ*m.

## Discussion

Although it is now well established that local protein synthesis is common to synapses in the adult brain [23, 42], the specificity of mRNA translation in different cell types and even in distinct connectivity motifs of the same neuron remains largely unexplored. Our work indicates that protein synthesis is regulated in a synapse-type specific manner during synapse formation. Tsc2, a key regulator of mTORC1 signaling in multiple cellular contexts [12], is critical for the development of excitatory synapses onto PV^+^ cells but not onto SST^+^ interneurons. This specificity is mediated by the activation of ErbB4, which controls excitatory synapse development through the inhibition of Tsc2 and the subsequent induction of a molecular program of mRNA translation involving the synthesis of several cell-adhesion and glutamate receptor-related proteins. While some of these molecules have been previously linked to glutamatergic synapses contacting interneurons [43–45], the involvement of most of them in the formation of these synapses was not previously known.

Our results reveal that local translation is directly involved in synapse formation. Recent work in *Drosophila* suggested that the local action of the phosphatase Prl-1 (phosphatase of regenerating liver 1) in synapse formation might be achieved by local translation [46]. Here, we found that translation occurs locally in developing synapses in the mammalian neocortex and that disrupting the machinery regulating protein synthesis deregulates synapse formation. Moreover, we demonstrate that many of the proteins identified as being synthesized at developing synapses are involved in synapse formation. Local translation, therefore, contributes to brain wiring not only during axon guidance [15-17, 47-49] but also in the final stages of neural circuit assembly.

Altered synthesis of synaptic proteins is a core pathophysiological mechanism in autism spectrum disorder (ASD) [50–55]. While the evidence linking *ERBB4* with intellectual disability and ASD is relatively scarce [56, 57], the identification of *NLGN3* as a target of ErbB4-Tsc2 signaling in the formation of excitatory synapses onto PV^+^ interneurons is intriguing because mutations in *TSC2* and *NLGN3* are strongly associated with ASD [58–62]. The excitation received by PV^+^ cells is strongly modulated during experience [41, 63, 64], which suggest a prominent role in learning and memory. The synapse-specific molecular program unveiled in this study reinforces the idea that this connection is a sensitive hub for maladaptive network responses in neurodevelopmental disorders.

## ACKNOWLEDGEMENTS

We thank T. Garcés and E. Serafeimidou-Pouliou for general laboratory support, I. Andrew for the management of mouse colonies, the Centre for Genomic Regulation (CRG) Genomics Unit for RNA-seq, and P. de la Grange at GenoSplice for help with bioinformatic analyses. We are also grateful to J. Bateman, M.J. Conde-Dusman, G. Condomitti, N. Flames, and C. Houart for critical reading of the manuscript, and members of the Marín and Rico laboratories for stimulating discussions and ideas. This work was supported by grants from the European Union’s Horizon 2020 Research and Innovation Programme AIMS-2-TRIALS (777394) and the Simons Foundation Autism Research Initiative (736666) to B.R. and O.M. C.B., D.E-A., M.S., B.R., and O.M. designed experiments. C.B. carried out the biochemical experiments and the histological analysis of *Tsc2* mutants. M.S. performed electrophysiological experiments. D.E-A. performed the functional analysis of target genes. A.H-G. and S.S. contributed to data collection and analysis. F.O. and P.M. produced the AAVs. C.B., B.R., and O.M. wrote the manuscript with input from all the authors.

## MATERIALS AND METHODS

### Mice

All experiments were performed following the guidelines of King’s College London Biological Service Unit and of the European Community Council Directive 86.609/EEC. Animal work was carried out under license from the UK Home Office following the Animals (Scientific Procedures) Act 1986. Animals were maintained under standard laboratory conditions on a 12:12 light/dark cycle with water and food ad libitum. Both male and female mice were used indiscriminately throughout the study. Mice carrying loxP-flanked *Tsc2* alleles [65] (JAX027458) were crossed with *Lhx6-Cre* mice [66] to generate the *Lhx6-Cre;Tsc2^F/F^* mouse line. F1 mice were crossed with *RCL^tdT^* mice [67] (JAX7909) to generate the *Lhx6-Cre;Tsc2^F/F^;RCL^tdT/tdT^* mice. Mice carrying loxP-flanked *Erbb4* alleles [68] were crossed with *Lhx6-Cre* mice to generate *Lhx6-Cre;Erbb4^F/F^* mutants. *Erbb4^F/F^* mice were crossed with *Tsc2^F/F^* mice to generate *Erbb4^F/F^;Tsc2^F/F^* mice, which were then crossed with *Lhx6-Cre;Erbb4^F/F^* mice to generate the *Lhx6-Cre;Erbb4^F/F^;Tsc2^F/+^* mice. *Erbb4^F/F^* mice were crossed with *Rpl22^HA/HA^* (RiboTag) mice [38] (JAX011029) to generate *Erbb4^F/F^;Rpl22^HA/HA^* mice, that were then crossed with *Lhx6-Cre;Erbb4^F/F^* mice to generate the *Lhx6-Cre;Erbb4^F/F^;Rpl22^HA/HA^* mice. C57BL/6 mice (Charles River) were used for biochemistry and single-molecule fluorescence in situ hybridization experiments, and CD1 mice (Crl:CD1[ICR], Charles River) were used for in utero electroporation.

### Histology

#### Immunohistochemistry

Mice were deeply anaesthetized with sodium pentobarbital by intraperitoneal injection and transcardially perfused with saline followed by 4% paraformaldehyde (PFA). Brains were postfixed for 2 h at 4 °C, cryoprotected in 15% sucrose followed by 30% sucrose and sectioned frozen on a sliding microtome (Leica SM2010R) at 40 *µ*m. Free-floating sections were permeabilized with 0.25% Triton X-100 in PBS for 1 h and blocked for 2 h in PBS containing 0.3% Triton X-100, 10% serum, and 5% BSA. Sections were then incubated overnight at 4 °C with primary antibodies. The next day, sections were washed in PBS and incubated with secondary antibodies for 2 h at room temperature. When required, sections were counterstained with 5 *µ*M DAPI in PBS. Sections were allowed to dry and mounted in Mowiol/DABCO. All primary and secondary antibodies were diluted in PBS with 0.3% Triton X-100, 5% serum, and 1% BSA. The following primary antibodies were used: guinea-pig anti-VGluT1 (1:2000, Chemicon #AB5905), mouse anti-PSD95 (1:500, NeuroMabs #75-028), chicken anti-PV (1:500, Synaptic Systems #195 006), mouse anti-PV (1:1000, Sigma-Aldrich #P3088), rat anti-SST (1:200, Millipore #MAB354), rabbit anti-dsRed (1:500, Clontech #632496), goat anti-mCherry (1:500, Antibodies Online #ABIN1440057), rabbit anti-P-S6rp (1:1000, Cell Signalling Technology #5364), mouse anti-Syt2 (1:250, ZFIN #ZDB-ATB-081002-25), mouse anti-Gephyrin (1:500, Synaptic Systems #147 011), rabbit anti-NeuN (1:500, Millipore #ABN78), rabbit anti-GluA4 (1:500, Millipore #AB1508), chicken anti-SynCAM (1:200, MBL International #CM004-3), goat anti-Nptn (1:500, Invitrogen #PA5-47726), rabbit anti-TrkC (1:500, Cell Signalling Technology #3376), rabbit anti-Cacng2 (1:100, Millipore #07-577), rabbit anti-Nlgn3 (1:400, Synaptic Systems #129 113), rabbit anti-Clstn2 (1:100, MyBioSource #MBS9208953). We used Alexa Fluor-conjugated (Invitrogen), DyLight-conjugated and Cy3-conjugated (Jackson ImmunoResearch) secondary antibodies. For biotin amplification, sections were incubated with biotinylated secondary antibody (1:200, Vector labs #BA-2000) followed by Alexa Fluor-conjugated Streptavidin (Invitrogen).

#### Single-molecule fluorescence in situ hybridization

All solutions for single-molecule fluorescent *in situ* hybridization were prepared in RNase-free PBS. Mice were perfused as described above and brains were postfixed overnight at 4°C, cryoprotected in 15% sucrose followed by 30% sucrose, and sectioned frozen on a sliding microtome at 30 *µ*m. Sections were mounted on RNase-free SuperFrost Plus slides (ThermoFisher) and probed against the candidate genes as well as *Parvalbumin* (*Pvalb*) using the RNAscope Multiplex Fluorescent Assay v2 protocol (ACDBio #323110). The following probes were used: Cadm1-C1 (Catalogue #492361), Nptn-C1 (Catalogue #1066021), Nlgn3-C1 (Catalogue #497661), Ntrk3-C1 (Catalogue #423621), Clstn2-C1 (Catalogue #542621), Gria4-C1 (Catalogue #422801), Cacng2-C1 (Catalogue #437221), Pvalb-C2 (Catalogue #421931).

### Synaptosome experiments

#### Synaptosomes preparation

Cortices from both brain hemispheres were rapidly dissected in ice-cold PBS and immediately homogenized in SynPER Synaptic Protein Extraction Reagent (ThermoScientific) complemented with cOmplete protease inhibitors (Sigma-Aldrich) and phosSTOP phosphatase inhibitors (Sigma-Aldrich). Synaptosomes were prepared according to the manufacturer’s instructions and processed for further applications. For Western blot, synaptosomes were denatured at 95 °C for 10 min in Laemmli sample buffer (80 mM Tris-HCl pH 6.8, 100 mM DTT, 8.7% glycerol, 2% SDS, and 0.01% bromophenol blue) and protein concentrations were measured using Pierce 660nm Protein Assay (ThermoFisher).

#### Co-immunoprecipitation

Cortical synaptosomes from P15 C57BL/6 mice were diluted in 1 ml of co-immunoprecipitation buffer (20 mM Tris-HCl pH 8, 120 mM NaCl, 1% NP40, 1% glycerol, and 1x cOmplete protease inhibitors) and subsequently incubated overnight at 4 °C with 5 *µ*g of one of the following antibodies: rabbit anti-Tsc2 (Cell Signalling Technology #4308), rabbit anti-ErbB4 (Santa Cruz #sc283) or rabbit IgG isotype control (Abcam #ab27478). 25 *µ*l of Dynabeads Protein G slurry (Invitrogen) was washed in co-immunoprecipitation buffer and added to the samples for 3 h at 4 °C with gentle rotation. Beads were washed six times with co-immunoprecipitation buffer, resuspended in Laemmli sample buffer 1x and co-immunoprecipitation samples were then denatured at 95 °C for 10 min.

#### Anti-HA pulldown

Cortical synaptosomes from P15 *Lhx6-Cre;Rpl22^HA/HA^* mice were diluted in 1 ml of lysis buffer (20 mM Tris-HCl pH 8, 120 mM NaCl, 1% NP40, 1% glycerol, 1x cOmplete protease inhibitors). 100 *µ*l of anti-HA magnetic beads (Pierce #88837) were washed in lysis buffer and incubated with the samples for 3-4 h at 4°C with gentle rotation. After incubation, beads were washed 6 times in ice-cold lysis buffer, resuspended in Laemmli sample buffer 1x and samples were then denatured at 95 °C for 10 min.

#### ErbB4 activation

Cortical synaptosomes were incubated at 35 °C with 450 rpm in PBS with 2 mM ATP (Sigma-Aldrich), 125 mM HEPES, 10 mM MgCl_2_, 67 *µ*M amino acid mixture (Promega), 1x cOmplete protease inhibitors EDTA-free, 10 *µ*M MG132 (Sigma-Aldrich), and phosSTOP phosphatase inhibitors. Synaptosomes were preincubated for 15 min with 50 *µ*M of cycloheximide (Sigma-Aldrich) (or DMSO for control) and then treated for 25 min with 500 ng/ml Nrg-EGF (Peprotech #100-03) or 0.1% BSA for control. The reaction was stopped by adding 1% SDS and the samples were denatured at 95 °C for 10 min in Laemmli sample buffer. Protein concentrations were measured using Pierce 660 nm Protein Assay.

#### Plating and immunofluorescence

Cortical synaptosomes were incubated for 90 min at 4 °C with gentle shaking on Nunc Lab-Tek 8-well chamber slides (Sigma-Aldrich) coated with poly-D-lysine (ThermoScientific). Synaptosomes were then fixed with 4% PFA for 15 min at room temperature and washed in PBS. Samples were permeabilized in PBS 0.5% Triton X-100 for 10 min, blocked with PBS 4% serum for 30 min at room temperature, and then incubated overnight at 4 °C with primary antibodies in PBS 4% serum. The next day, samples were washed in PBS and incubated with secondary antibodies for 2 h at room temperature. The samples were then washed in PBS and mounted in Mowiol/DABCO. The following primary antibodies were used: guinea-pig anti-VGluT1 (1:2000, Chemicon #AB5905), mouse anti-PSD95 (1:500, NeuroMabs #75-028), rabbit anti-P-S6rp (1:1000, Cell Signalling Technology #5364) and goat anti-Nrg3 (1:500, Neuromics #GT15220). We used Alexa Fluor-conjugated (Invitrogen) and Cy3-conjugated (Jackson ImmunoResearch) secondary antibodies.

### Imaging and analysis

#### Image acquisition

Images were acquired with an inverted SP8 confocal microscope (Leica) or an ApoTome (Zeiss), using the LAS AF software and the ApoTome function in Zen2 software, respectively. Samples from the same experiment were imaged and analyzed in parallel, using the same laser power, photomultiplier gain and detection filter settings. Imaging of synaptic markers and in situ hybridization was performed at 8-bit depth, with 100x objective and 2.2 digital zoom at 200 Hz acquisition speed. Imaging of P-S6rp intensity in cortical tissue and synaptosomes was performed at 12-bit depth, with a 40x objective and 200 Hz of acquisition speed, or a 63x objective, 2.2 digital zoom and 200 Hz of acquisition speed, respectively. Imaging of post-recording immunostaining was performed at 8-bit depth, with a 20x objective at 200Hz acquisition speed.

#### Image analysis

Analysis of cell volume and P-S6rp intensity was performed using Imaris 8.1.2 (Bitplane). Background subtraction and Gaussian filtering were first applied in all channels. Cell somas were reconstructed automatically as three-dimensional isosurfaces with the “create surface” tool using the PV staining (for PV^+^ interneurons) or the tdTomato staining (for SST^+^ interneurons). The volume of the reconstructed somas was then quantified automatically as well as the intensity of the P-S6rp channel (as integrated density corrected for soma size) inside each reconstructed soma.

Analysis of cluster/synaptic densities was performed using a custom macro in FIJI (ImageJ), as previously described [37, 41]. Background subtraction, Gaussian blurring, smoothing, and contrast enhancement were first applied in all channels. Cell somas were drawn automatically or manually based on intensity levels of PV staining (for PV^+^ interneurons), tdTomato staining (for SST^+^ interneurons) or NeuN staining (for pyramidal cells), to create a mask of the soma surface and measure its perimeter. Presynaptic boutons and postsynaptic clusters were detected automatically based on thresholds of intensity (thresholds for the different synaptic markers were selected from a set of random images before quantification and the same threshold was applied to all images from the same experiment). The “Analyze Particles” and “Adjustable Watershed” tools were applied to the synaptic channels and a mask was generated with a minimum particle size of 0.05. The soma and synaptic masks were merged to automatically quantify the number of puncta contacting the soma. Puncta were defined as presynaptic boutons contacting the soma surface when ≥ 0.04 *µ*m^2^ of the puncta area in the synaptic mask was colocalizing with the soma mask. Puncta were defined as postsynaptic clusters contained inside a soma when ≥ 0.04 *µ*m^2^ of the puncta area in the synaptic mask was colocalizing with the soma mask. Synapses were defined when a presynaptic bouton and a postsynaptic cluster were contacting each other with a colocalisation area of ≥ 0.03µm^2^ of their corresponding masks.

Analysis of P-S6rp intensity in synaptosomes was performed using a custom macro in FIJI. Background subtraction, Gaussian blurring, smoothing, and contrast enhancement were first applied in all channels. Pre- and postsynaptic puncta were detected automatically based on thresholds of intensity (thresholds for the different synaptic markers were selected from a set of random images before quantification and the same threshold was applied to all images from the same experiment). The “Analyze Particles” and “Adjustable Watershed” tools were applied to the synaptic channels and a mask was generated with a minimum particle size of 0.05. The two synaptic masks were merged to automatically quantify the number of synaptosomes, with a synaptosome identified when pre- and postsynaptic puncta were contacting each other with a colocalization area of ≥ 0.03 *µ*m^2^ of their corresponding masks. The intensity of the P-S6rp channel was automatically calculated in each defined synaptosome.

Analysis of in situ hybridization was performed using a custom macro in FIJI. Background subtraction, Gaussian blurring, smoothing, and contrast enhancement were first applied in all channels. Cell somas were drawn automatically or manually based on intensity levels of *Pvalb* in situ hybridization signal, to create a mask of the soma surface and measure its perimeter. RNA particles were detected automatically based on thresholds of intensity (thresholds for the different target RNA were selected from a set of random images before quantification and the same threshold was applied to all images from the same experiment). The “Analyze Particles” and “Adjustable Watershed” tools were applied to the target RNA channels and a mask was generated with a minimum particle size of 0.05. The soma and target RNA masks were merged to automatically quantify the number of particles inside the soma.

Analysis of target staining intensity was performed using a custom macro in FIJI. Background subtraction, Gaussian blurring, smoothing, and contrast enhancement were first applied in all channels, Cell somas were drawn manually to create a mask of the soma surface. The intensity was measured automatically as raw integrated density.

### Western blotting

10-40 *µ*g of denatured protein extracts were separated by SDS-PAGE using 10% acrylamide gels or 4-15% Mini-Protean precast gels (Bio-Rad) for 2 h at 120 V and transferred onto methanol-activated PVDF membranes at 350 mA for 2 h on ice. Membranes were blocked with either 5% non-fat milk or 5% BSA in TBS-T (20mM Tris-HCl pH 7.5, 150mM NaCl, and 0.1% Tween20) for 1 h at room temperature. Membranes were then probed with primary antibodies in corresponding blocking buffer overnight at 4 °C. The next day, membranes were washed in TBS-T and incubated with HRP-conjugated secondary antibodies for 1 h at room temperature. Depending on the amount of protein of interest in the samples, membranes were incubated with either Immobilon Western Chemiluminescent HRP substrate (Millipore), SuperSignal West Pico or West Femto Chemiluminescent Substrates (ThermoFisher). Protein levels were visualized by chemiluminescence with an Odyssey FC (Li-Cor) and quantified with Image Studio Lite. For quantification, densitometry of the band of interest was normalized to that of actin. For phosphorylated proteins, densitometry of the band of interest was normalized to that of the corresponding total protein.

The following primary antibodies were used: rabbit anti-ErbB4 (1:1000, Cell Signalling Technology #4795), rabbit anti-P-S6rp (Ser240/244, 1:5000, Cell Signalling Technology #5364), rabbit anti-S6rp (1:5000, Cell Signalling Technology #2217), rabbit anti-P-Tsc2 (Thr1462, 1:1000, Cell Signalling Technology #3617), rabbit anti-Tsc2 (1:1000, Cell Signalling Technology #4308), rabbit anti-P-4EBP1 (Thr37/46, 1:1000, Cell Signalling Technology #2855), rabbit anti-4EBP1 (1:1000, Cell Signalling Technology #9644), mouse anti-actin-peroxidase (1:20 000, Sigma-Aldrich #A3854), mouse anti-HA (1:500, ThermoFisher #26183), rabbit anti-GluA4 (1:1000, Millipore #AB1508), chicken anti-SynCAM (1:1000, MBL International #CM004-3), goat anti-Nptn (1:1000, Invitrogen #PA5-47726), rabbit anti-TrkC (1:1000, Cell Signalling Technology #3376), rabbit anti-Cacng2 (1:1000, Millipore #07-577), rabbit anti-Nlgn3 (1:1000, Synaptic Systems #129 113) and rabbit anti-Clstn2 (1:1000, MyBioSource #MBS9208953). The following HRP-conjugated secondary antibodies were used: goat anti-mouse-peroxidase (1:5000, Invitrogen #31444), donkey anti-rabbit-peroxidase (1:5000, Invitrogen #SA1-200), donkey anti-chicken-peroxidase (1:5000, Invitrogen #SA1-300), rabbit anti-goat-peroxidase (1:5000, Abcam #ab6741). For immunoprecipitation samples, the following secondary antibodies were used: anti-mouse-HRP (1:1000, Abcam #ab131368) and anti-rabbit-HRP (1:1000, Abcam #ab131366).

### In vitro electrophysiology

#### Slice preparation and patch-clamp recordings

Mice were anaesthetized with an overdose of sodium pentobarbital before decapitation. Coronal slices of 300 μm were cut using a VT1200S vibratome (Leica) in ice-cold artificial cerebrospinal fluid (ACSF) containing 87 mM NaCl, 11 mM glucose, 75 mM sucrose, 2.5 mM KCl, 1.25 mM NaH_2_PO_4_, 0.5 mM CaCl_2_, 7 mM MgCl_2_, 26 mM NaHCO_3_, oxygenated with 95% O2 and 5% CO2, and incubated for 1 h at 32 °C and subsequently at room temperature. Slices were transferred to the recording setup 15 min prior to recording and incubated at 32 °C while being continuously oxygenated with 95% O2 and 5% CO2 in recording ACSF containing: 124 mM NaCl, 1.25 mM NaH_2_PO_4_, 3 mM KCl, 26 mM NaHCO_3_, 10 mM Glucose, 2 mM CaCl_2_, 1 mM MgCl_2_. Pipettes (3–5 MΩ) were made from borosilicate glass capillaries using a PC-10 pipette puller (Narishige) and filled with intracellular solution containing 130 mM potassium-gluconate, 5 mM KCl, 10 mM HEPES, 2.5 mM MgCl_2_, 4mM Na2ATP, 0.4mM Na3GTP, 10 mM sodium-phosphocreatine, 0.6 mM EGTA (pH 7.2–7.3, 285–295mOsm) supplemented with either 0.2 mg/ml neurobiotin (Vector Laboratories) for current clamp recordings, or 115 mM CsMeSO_3_, 20 mM CsCl, 10 mM HEPES, 2.5 mM MgCl_2_, 4 mM Na2ATP, 0.4 mM Na3GTP, 10 mM sodium-phosphocreatine, 0.6 mM EGTA (pH 7.2–7.3, 285–295 mOsm), 0.2% neurobiotin for voltage clamp recordings. Traces were recorded using a Multiclamp 700B amplifier (Molecular Devices), sampled at 50 kHz and filtered at 3 kHz. sEPSCs were recorded in the presence of 100 *µ*M Picrotoxin (Tocris Bioscience) at a holding potential of −60mV and analyzed using Mini Analysis (Synaptosoft). Intrinsic properties were analyzed using Clampfit 10.2.

### Data analysis

Intrinsic properties were defined and analyzed as follows: RMP, the membrane potential recorded immediately after entering whole-cell configuration; TC, fitting of a single exponential to the response to a 10 mV hyperpolarizing step; IR, an average of the resistance calculated from 5 increasing 10 pA current steps; Ih sag, calculated from the average membrane potential at the end of a 1s current pulse, initially hyperpolarizing the cell to –100 mV; APmax, the maximum value reached by the AP; MFF, Maximum AP frequency elicited by increasing 100pA steps; fAHP, the potential difference between the threshold and minimum AP-value; and Capacitance, calculated from TC and IR.

#### Post-recording immunohistochemistry

After recording, slices were immediately fixed in PFA 4% for 30 min at 4 °C washed in PBS and permeabilized with PBS containing 0.4% Triton X-100 for 1 h at room temperature. Slices were blocked in PBS with 0.3% Triton X-100, 10% serum, and 5% BSA for 3 h at room temperature and incubated with primary antibodies overnight at 4 °C. Slices were then washed in PBS and incubated with secondary antibodies overnight at 4°C. The next day, slices were washed in PBS, counterstained with 5 *µ*M DAPI in PBS, and mounted in Mowiol/DABCO. All primary and secondary antibodies were diluted in PBS containing 0.3% Triton X-100, 5% serum, and 1% BSA. The following primary and secondary antibodies were used: rat anti-SST (1:200, Millipore #MAB354), chicken anti-PV (1:500, Synaptic Systems #195 006) rabbit anti-dsRed (1:500, Clontech #632496), goat anti-rat 647 (1:400, Invitrogen #A-21247), donkey anti-chicken 405 (1:200, Jackson ImmunoResearch #703-475-155) and donkey anti-rabbit Cy3 (1:500, Jackson ImmunoResearch #711-165-152). Neurobiotin was revealed with Streptavidin 488 (1:400 Invitrogen #S11223).

### RNA isolation by Ribotrap pulldown

Cortices from P15 *Lhx6-Cre;Erbb4^+/+^;Rpl22^HA/HA^* and *Lhx6-Cre;Erbb4^F/F^;Rpl22^HA/HA^* mice were rapidly dissected in ice-cold RNase-free PBS and immediately homogenized in SynPER Synaptic Protein Extraction Reagent (ThermoScientific) supplemented with 1x cOmplete EDTA-free protease inhibitors (Sigma-Aldrich), 1 mg/ml Heparin (Sigma-Aldrich), 200 U/ml RNAsin (Promega), 100 *µ*g/ml cycloheximide and 1 mM DTT (Sigma-Aldrich). Synaptosomes were then prepared according to the manufacturer’s instructions. The final synaptosome pellets were resuspended in supplemented SynPER and Igepal-CA630 (Sigma-Aldrich) was added to the samples to a final concentration of 1%. 100 *µ*l of anti-HA magnetic beads (Pierce #88837) were washed in supplemented SynPER and added to the samples for 3-4 h at 4°C with gentle rotation. After incubation, beads were washed 3 times in ice-cold washing buffer (300 mM KCl, 1% Igepal-CA630, 50 mM Tris-HCl pH 7.5, 12mM MgCl_2_, 1mM DTT and 100 *µ*g/ml cycloheximide) and eluted in 350 *µ*l of RLT Plus buffer from the RNAeasy Plus Micro kit (Qiagen) supplemented with 2-mercaptoethanol (Bio-Rad).

### RNA sequencing and differential expression analysis

#### RNA purification, quantification, and quality check

RNA purification of immunoprecipitated RNA was performed using the RNeasy Plus Micro kit (Qiagen) following the manufacturer’s protocol. RNA quality was checked on a Bioanalyzer instrument (Agilent Technologies) using an RNA 6000 Pico Chip. Only RNA samples with RNA integrity number (RIN) values higher than 7.4 were used for library preparation and sequencing.

#### Library preparation and Illumina sequencing

Four biological replicates were analyzed for each genotype. Library preparation and RNA sequencing experiments were performed by the Genomic Unit of the Centre for Genomic Regulation (CRG, Barcelona, Spain). Library preparation for all samples was performed with 1 ng of total RNA using the SMARTer Ultra Low RNAkit for ultra-low RNA. Samples were then sequenced paired-end using an Illumina HiSeq 2500 platform to a mean of approximately 60-80 million mapped reads per sample.

#### Quality control and differential gene expression analysis

RNA sequencing data analysis was performed by GenoSplice (www.genosplice.com). Sequencing, data analysis, reads repartition, and insert size estimation were performed using FastQC, Picard-Tools, Samtools and rseqc. Reads were mapped using STAR v2.4.0 [69] on the mm10 Mouse genome assembly with an average of 79.8±3.4% of uniquely mapped reads. The analysis of gene expression regulation was performed as described previously [70]. In brief, for each gene present in the Mouse Fast DB v2021_1 annotation, reads aligning on constitutive regions (that are not prone to alternative splicing) were counted. Based on these read counts, normalization and differential gene expression were performed using DESeq2, as described before [71], on R v3.2.5. Genes were considered as expressed if their Reads Per Kilobase of transcript per Million mapped reads (RPKM) value was greater than 99% of the background Fragments Per Kilobase of transcript per Million mapped reads (FPKM) value based on intergenic regions. Only genes expressed in at least one of the two compared experimental conditions were further analyzed. Results were considered statistically significant for p-values ≤ 0.05 and fold changes ≥ 1.5. Pathway and GO analyses were performed using WebGestalt [72] and were considered significant for p-values ≤ 0.05.

#### Candidate selection criteria

From the list of differentially expressed genes downregulated in *Erbb4* conditional mutants compared to controls, we used the Synaptic Gene Ontology tool (SynGO, https://syngoportal.org), to highlight genes coding for proteins with synaptic localization and/or function [39]. Among the “postsynapse” cellular component annotation, the terms “postsynaptic specialization” and “postsynaptic membrane” were significantly enriched when using the following settings: stringency = remove annotations with only proteomics evidence; minimum gene counts per term = 5. Merging both lists, we removed genes that were not expressed in PV^+^ interneurons at P10 (FPKM < 5 in Synapdomain [7]; https://devneuro.org/cdn/synapdomain.php). The resulting list of genes was classified into further subcategories according to the gene ontology analysis or a manual MEDLINE search. The two subcategories with the highest number of genes were “cell adhesion molecules” and “AMPA receptor-related molecules”. We used a set of four additional criteria to select candidates for functional validation: (1) expression in PV^+^ interneurons at P10 (Synapdomain) [7], (2) expression in PV^+^ interneurons at P30 (SpliceCode) [73], available at https://scheiffele-splice.scicore.unibas.ch, (3) enrichment in PV^+^ versus SST^+^ interneurons (SpliceCode), and (4) literature on interneuron connectivity (MEDLINE search for “gene name” and “synapse” and “interneuron”). Each of these criteria was given a score and the sum of the four scores was used to rank the genes. In the “cell adhesion molecules” category, 8 genes had 3 or more points. Out of these 8 genes, we found three calsyntenin-encoding genes and two neuroligin-encoding genes. To avoid validating genes belonging to the same protein family, we selected *Clstn2* and *Nlgn3* for functional analyses. The other three selected genes were *Nptn*, *Cadm1*, and *Ntrk3*. In the “AMPA receptor-related molecules” category, the two top-ranking genes were *Gria4* and *Cacng2*.

### AAV experiments

#### Generation of AAV expression vectors

To downregulate the expression of the target genes *in vivo*, we took advantage of a cell type-specific knockdown strategy using a Cre-dependent cassette *pDIO-DSE-mCherry-PSE-MCS* as previously reported [7]. We generated five different *shRNA* constructs targeting each of the target genes using the Block-iT RNAi Designer web tool (Thermo Scientific) to identify target sequences within the coding region of the genes that show high knockdown effectivity *in silico*. The target sequences for each *shRNA* construct were the following: *shLacZ* (AAA TCG CTG ATT TGT GTA GTC), *shCadm1* (shRNA-1: GGG AGG AGA TTG AAG TCA ACT; shRNA-2: GGT TCA AAG GGA ACA AGG; shRNA-3: GCA GTA TAA ACC GCA AGT GCA; shRNA-4: GCA TTT GAG TTA ACG TGT GAA; shRNA-5: GGC CAA ACC TGT TCA ATA), *shNptn* (shRNA-1: GCT GAG GAT TCA GGC GAA TAC; shRNA-2: GCA GGA TGC TAT GAT GTA CTG; shRNA-3: GGT GTG TTT GAG ATT TCT; shRNA-4: GGC TGA AAT CAT CCT TGT; shRNA-5: GGT GAT CAT TGT GTA TGA), *shNlgn3* (shRNA-1: GCG AGG ACT TAG CGG ATA ATG; shRNA-2: GGA TAT GGT GGA TTG TCT TCG; shRNA-3: GCT ATG GCT CAC CTA CCT ACT; shRNA-4: GCA TGA CAT GTT CCA CTA TAC; shRNA-5: CAC CAT CAC TAT GAT TCC TAA), *shNtrk3* (shRNA-1: GCC AGA GCC TTT ACT GCA TCA; shRNA-2: GGA CCA ATG TAC ATG CCA TCA; shRNA-3: GGA ACA TTG CAT TGA GTT TGT; shRNA-4: GCA CAG ATT TCT TTG ACT TTG; shRNA-5: GGC ATC ACT ACA CCA TCA TCG), *shClstn2* (shRNA-1: GCA TCA CTA TGC CCT GTA TGT; shRNA-2: GGA GCA ACA TAT GAA CCA TAC; shRNA-3: GGA TAA AGT ATC ACT TCA ACC; shRNA-4: GGA CTT GGA TCC AAG GCA AGA; shRNA-5: GCA GGA GTC ATA AAC ATT TGG), *shGria4* (shRNA-1: GCA ACT AGA GCT TGA; shRNA-2: GCA CGT CAA AGG CTA CCA TTA; shRNA-3: GGA TCT GAA ACA CCT CCA AAG; shRNA-4: GGA ATT GAC ATG GAG AGA ACA; shRNA-5: GCA ATG ACA CAG CTA TCG), *shCacng2* (shRNA-1: GCC TCG AAG GGA ACT TCA AAG; shRNA-2: GCA AGC AAA TCG ACC ACT TTC; shRNA-3: GCT GAC ACC GCA GAG TAT TTC; shRNA-4: GCT GGC CGT GCA CAT GTT TAT; shRNA-5: GGA CAG GGA TAA CAG CTT TCT). For *in vitro* validation of knockdown effectivity of the *shRNA* constructs, the full-length coding regions of the target genes tagged with the hemagglutinin (HA)-tag (TAC CCC TAC GAC GTG CCC GAC TAC GCC) were designed and cloned in the pcDNA3.1(+) expression vector using GeneArt Gene Synthesis Services (Thermo Fisher). The reference coding sequences and the HA-tag insertion sites for the different target genes were as follows: HA-Cadm1 (NM_207676.2; HA-tag inserted after amino acid 420, alanine), HA-Nptn (NM_001357751.1; HA-tag inserted after amino acid 393, asparagine), HA-Nlgn3 (NM_172932.4; HA-tag inserted after amino acid 34, threonine), HA-Ntrk3 (NM_008746.5; HA-tag inserted after amino acid 1, methionine), HA-Clstn2 (NM_022319.2; HA-tag inserted after amino acid 22, glycine), HA-Gria4 (NM_019691.5; HA-tag inserted after amino acid 3, isoleucine), HA-Cacng2 (NM_007583.2; HA-tag inserted after amino acid 105, serine).

#### Cell culture and transfection

HEK293T and COS-7 cells were cultured in Dulbecco’s Modified Eagle’s medium supplemented with 10% fetal bovine serum, 2 mM glutamine, penicillin (50 units/ml) and streptomycin (50 g/ml). The cultures were incubated at 37 °C in a humidified atmosphere containing 5% CO_2_. Cells were transfected using FuGENE transfection reagent (ThermoFisher) at a 3:1 DNA:FuGENE ratio. Cells were seeded on 12-well plates in triplicate experiments and co-transfected with an *shRNA*-expressing construct, the expression plasmid for the corresponding HA-tagged gene, and a plasmid for expression of Cre recombinase. Co-transfection with a *shLacZ* construct was performed as a control for each target gene. 72 h after transfection, cells were scraped in RIPA lysis buffer containing 50 mM Tris-HCl pH 8, 150 mM NaCl, 0.5% sodium deoxycholate, 1% NP40, 0.1% SDS and 1x cOmplete protease inhibitors. Samples were denatured at 95 °C for 10 min in Laemmli sample buffer and protein concentrations were measured using Pierce 660 nm Protein Assay.

#### AAV production

HEK293T cells were seeded on 15-cm plates and co-transfected with packaging plasmids AAV-ITR-2 genomic vectors (7.5 *µ*g), AAV-Cap8 vector pDP8 (30 *µ*g, PlasmidFactory GmbH, Germany, #pF478) using PEI (Sigma) at a 1:4 DNA:PEI ratio. 72 h post-transfection, supernatants were incubated with ammonium sulphate (65 g/200 ml supernatant) for 30 min on ice and centrifuged at 4000 rpm for 45 min at 4 °C. Transfected cells were harvested and lysed (150 mM NaCl, 50 mM Tris pH 8.5), followed by three freeze-thaw cycles and benzonase treatment (50U/mL; Sigma) for 1 h at 37 °C. Filtered AAVs (0.8 *µ*m and 0.45 *µ*m MCE filters) from supernatants and lysates were run on an iodixanol gradient by ultracentrifugation (Vti50 roto, Beckmann Coulter) at 50 000 rpm for 1 h at 12 °C. The 40% iodixanol fraction containing AAVs was collected, concentrated using 100 kDa-MWCO Centricon plus-20 and Centricon plus-2 (Merck-Millipore), aliquoted and stored at -80 °C. The number of genomic copies was determined by quantitative real-time PCR using the following primers targeting the WPRE sequence (Fw: GGC ACT GAC AAT TCC GTG GT, and Rv: CGC TGG ATT GAG GGC CGA A). After *in vitro* validation, the following shRNA constructs were selected for AAV production based on their high knockdown efficiency: shCadm1 (shRNA-1 and shRNA-5), shNptn (shRNA-1 and shRNA-3), shNlgn3 (shRNA-2), shNtrk3 (shRNA-3 and shRNA-4), shClstn2 (shRNA-1 and shRNA-4), shGria4 (shRNA-2), and shCacng2 (shRNA-2 and shRNA-4).

### Surgeries

#### Stereotaxic injections

For intracranial injections of *shRNA*-expressing AAV vectors, P2-3 *Lhx6-Cre* pups were anaesthetized with isoflurane and mounted on a stereotactic frame using a 3D printed isoflurane mask. To target MGE-derived interneurons in layer 2/3 of the somatosensory cortex, the following coordinates were used: anteroposterior +1.6/1.7 mm; mediolateral -1.8/-1.9 mm; depth -0.3 mm.

#### In utero electroporation

In utero electroporation was performed as previously described [37]. Timed-pregnant CD1 females were deeply anaesthetized with isoflurane. Buprenorphine (Vetergesic, Ceva Animal Health Ltd) was administered for analgesia via subcutaneous injection, and ritodrine hydrochloride (Sigma-Aldrich #R0758) was applied to the exposed uterine horns to relax the myometrium. The DNA solution was mixed with Fast Green (Roche #06402612001) and 1-2 *µ*l of the solution was injected into the lateral ventricle of embryonic day 14.5 embryos. To target DNA into cortical pyramidal cell progenitors of the subventricular zone, five electric pulses (45 V for 50 ms, with 950 ms intervals) were delivered through electrodes (CUY650P3, Nepa Gene) connected to an electroporator (NEPA21 Super Electroporator, Nepa Gene).

### Statistical analyses

All statistical analyses were performed using Prism 9 (GraphPad Software). In general, no statistical methods were used to predetermine sample sizes and chosen samples size were like those reported in previous publications or generally employed in the field. Biological replicates (n values are different samples derived from different brains from different litters) were analyzed to assess the biological variability and reproducibility of data. To obtain unbiased data, experimental mice from all genotypes or conditions were processed together. Samples were tested for normality using the Shapiro–Wilk normality test. Unless otherwise stated, data were analyzed by t-test or ANOVA followed by post hoc Tukey’s analysis for comparisons of multiple samples. Differences were considered significant when p values < 0.05. Data are presented as mean ± s.e.m. Statistical details of experiments are described in figure legends.

---

**Suppl. Fig. 1.**
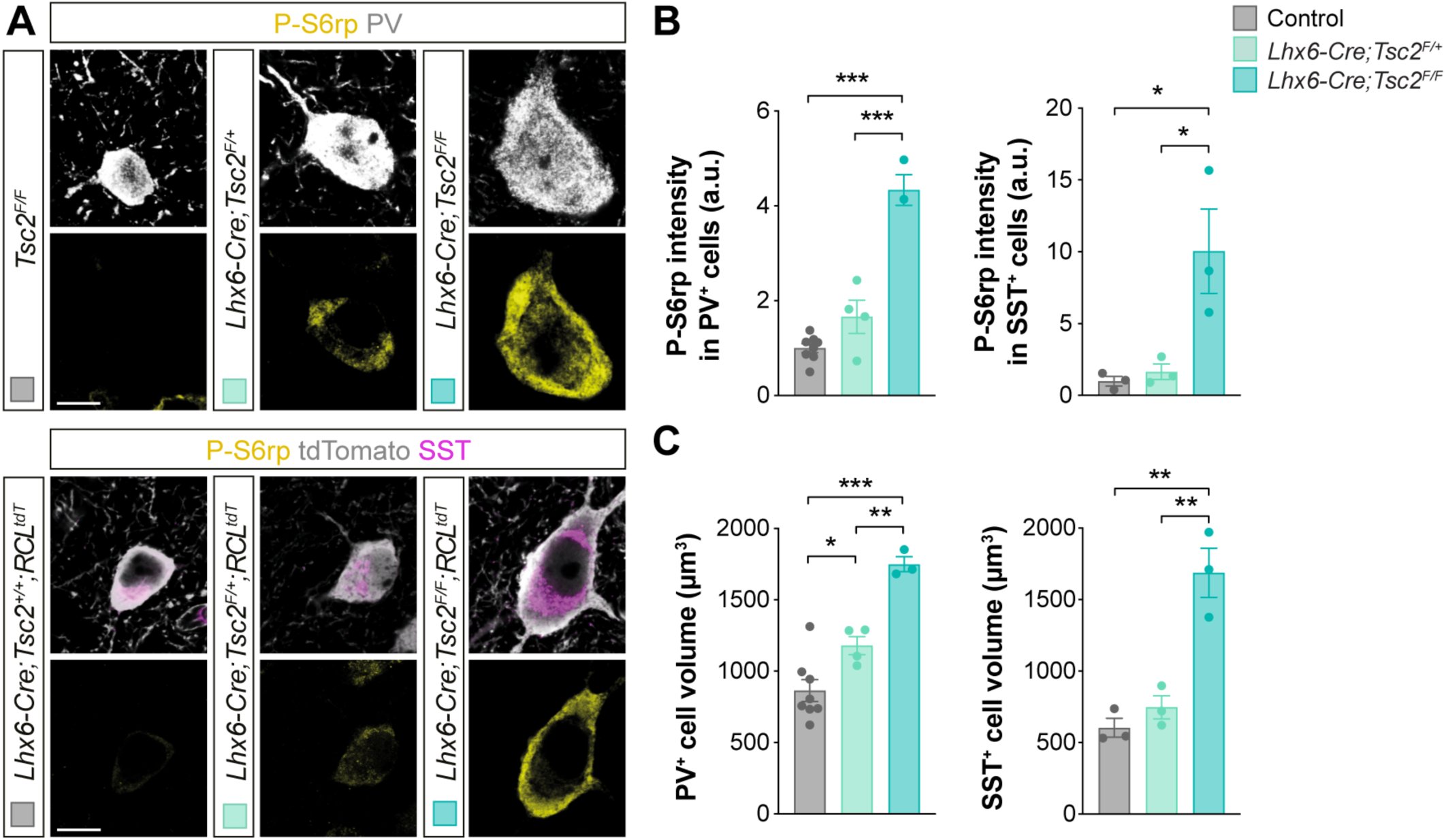
MGE-derived interneurons exhibit abnormally increased mTOR activation in conditional *Tsc2* mutants. (**A**), Confocal images illustrating phosphorylation of S6rp (P-S6rp, yellow) in PV^+^ interneurons (grey) (top) and SST^+^ (magenta) tdTomato^+^ (grey) interneurons (bottom) from P18-21 control, heterozygous and homozygous conditional *Tsc2* mutants. (**B**) Quantification of P-S6rp staining intensity in PV^+^ (left) and SST^+^ (right) interneurons in control, heterozygous and homozygous conditional *Tsc2* mutants. (**C**) Quantification of cell volume in PV^+^ (left) and SST^+^ (right) interneurons in control, heterozygous and homozygous conditional *Tsc2* mutants. One-way ANOVA followed by Tukey’s multiple comparisons test: *P < 0.05, **P < 0.01, ***P < 0.001 (PV^+^ cells: control *n* = 313 cells from 8 mice; heterozygous*, n* =125 cells from 4 mice; homozygous*, n* = 104 cells from 3 mice. SST^+^ cells: control, *n* = 61 cells from 3 mice; heterozygous*, n* =75 cells from 3 mice; homozygous*, n* = 61 cells from 3 mice). Data are mean ± s.e.m. Scale bar, 10 *µ*m.

**Suppl. Fig. 2.**
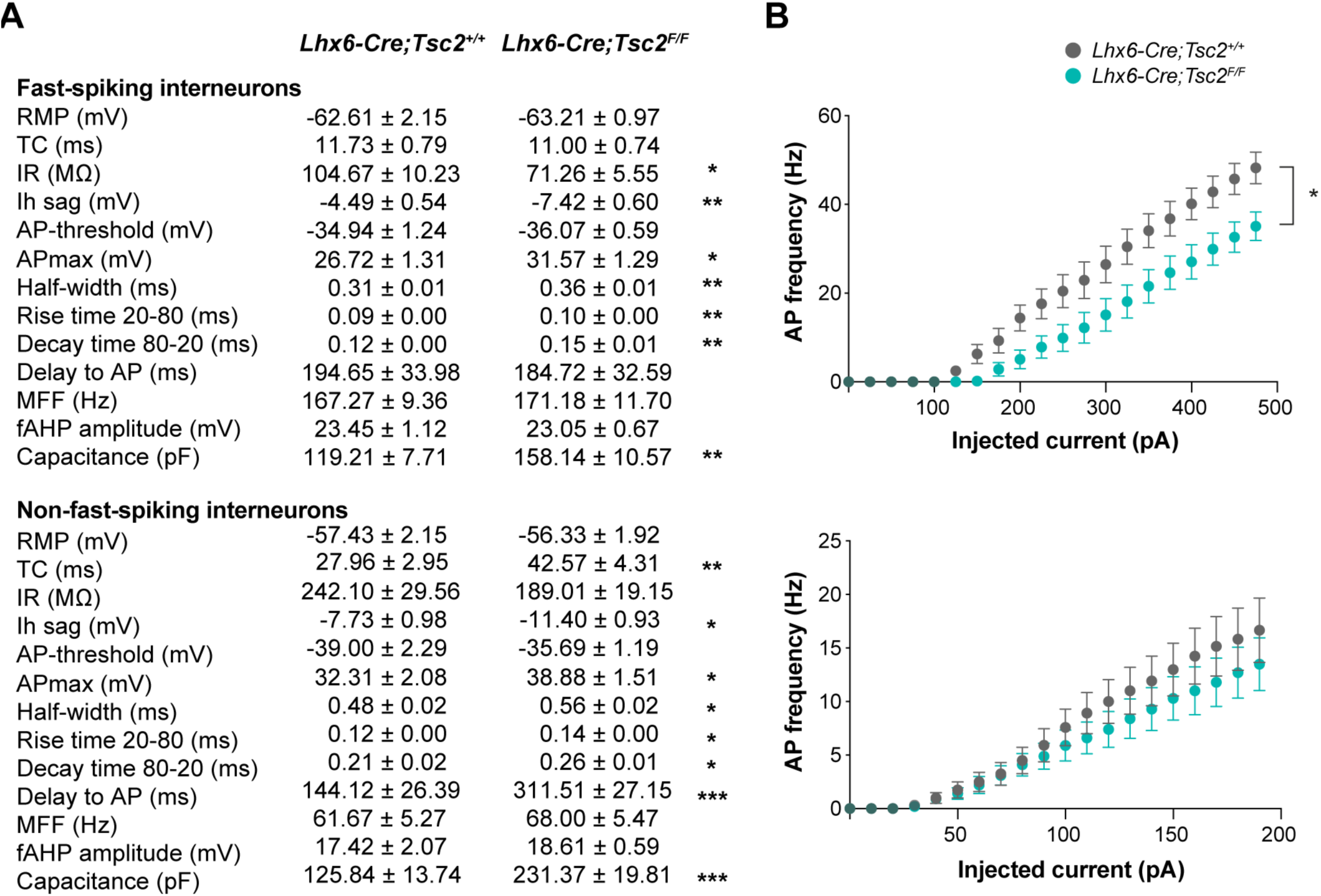
Intrinsic properties of MGE-derived interneurons in conditional *Tsc2* mutants. (**A**) Intrinsic electrophysiological properties of fast-spiking (top, putative PV^+^ interneurons) and non-fast-spiking (bottom, putative SST^+^ interneurons) interneurons from P18-21 control and homozygous conditional *Tsc2* mutants. RMP: resting membrane potential, TC: time constant, IR: input resistance, AP: action potential, MFF: maximum firing frequency, fAHP: fast afterhyperpolarization. (**B**) Excitability curves showing the spike frequency of fast-spiking (top) and non-fast-spiking (bottom) interneurons from P21 control and homozygous conditional *Tsc2* mutants in response to current injections. Two-way ANOVA with repeated measures: *P < 0.05 (fast-spiking interneurons: control, *n* = 15 cells from 8 mice; homozygous, *n* = 11 cells from 4 mice. Non-fast-spiking interneurons: control, *n* = 12 cells from 6 mice; homozygous, *n* = 10 cells from 4 mice). Data are mean ± s.e.m.

**Suppl. Fig. 3.**
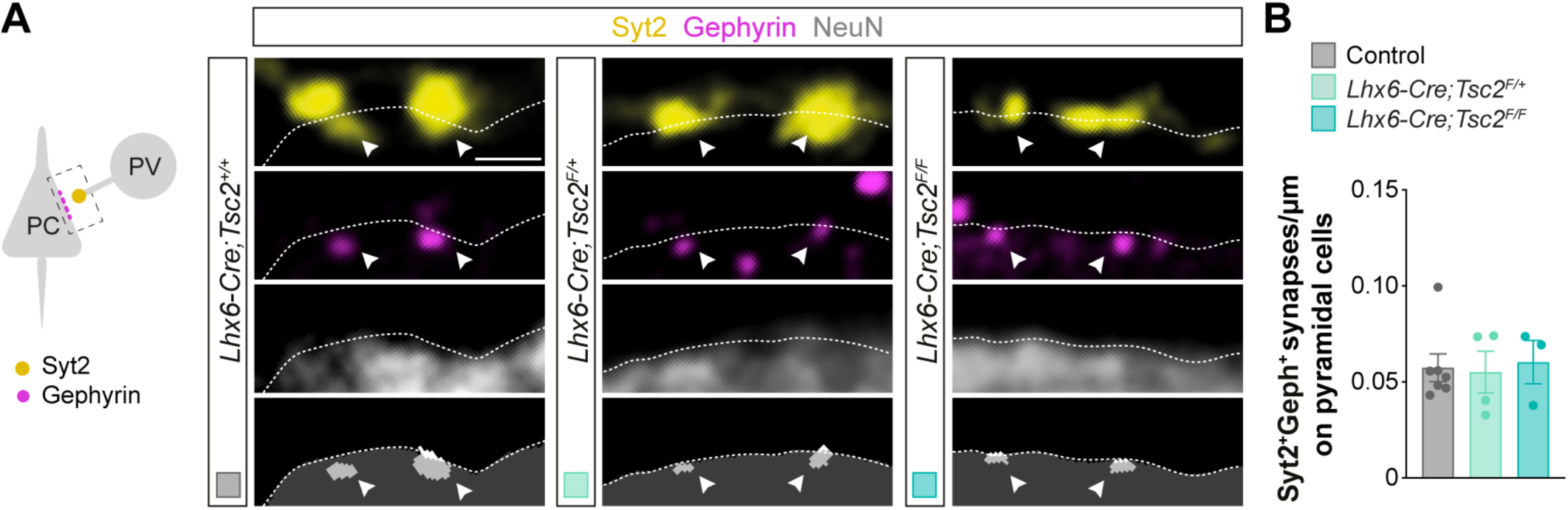
*Tsc2* deletion does not affect the synaptic output of PV^+^ basket interneurons. (**A**) Schematic of synaptic markers analyzed (left). Confocal images (top) and binary images (bottom) illustrating presynaptic Syt2^+^ puncta (yellow) and postsynaptic Gephyrin^+^ clusters (magenta) in NeuN^+^ pyramidal cells (grey) from P18-21 control, heterozygous and homozygous conditional *Tsc2* mutants. (**B**) Quantification of the density of Syt2^+^Gephyrin^+^ synapses contacting pyramidal cells (control, *n* = 112 cells from 7 mice; heterozygous*, n* = 71 cells from 4 mice; homozygous*, n* = 36 cells from 3 mice). One-way ANOVA followed by Tukey’s multiple comparisons test. Data are mean ± s.e.m. Scale bar, 1 *µ*m.

**Suppl. Fig. 4.**
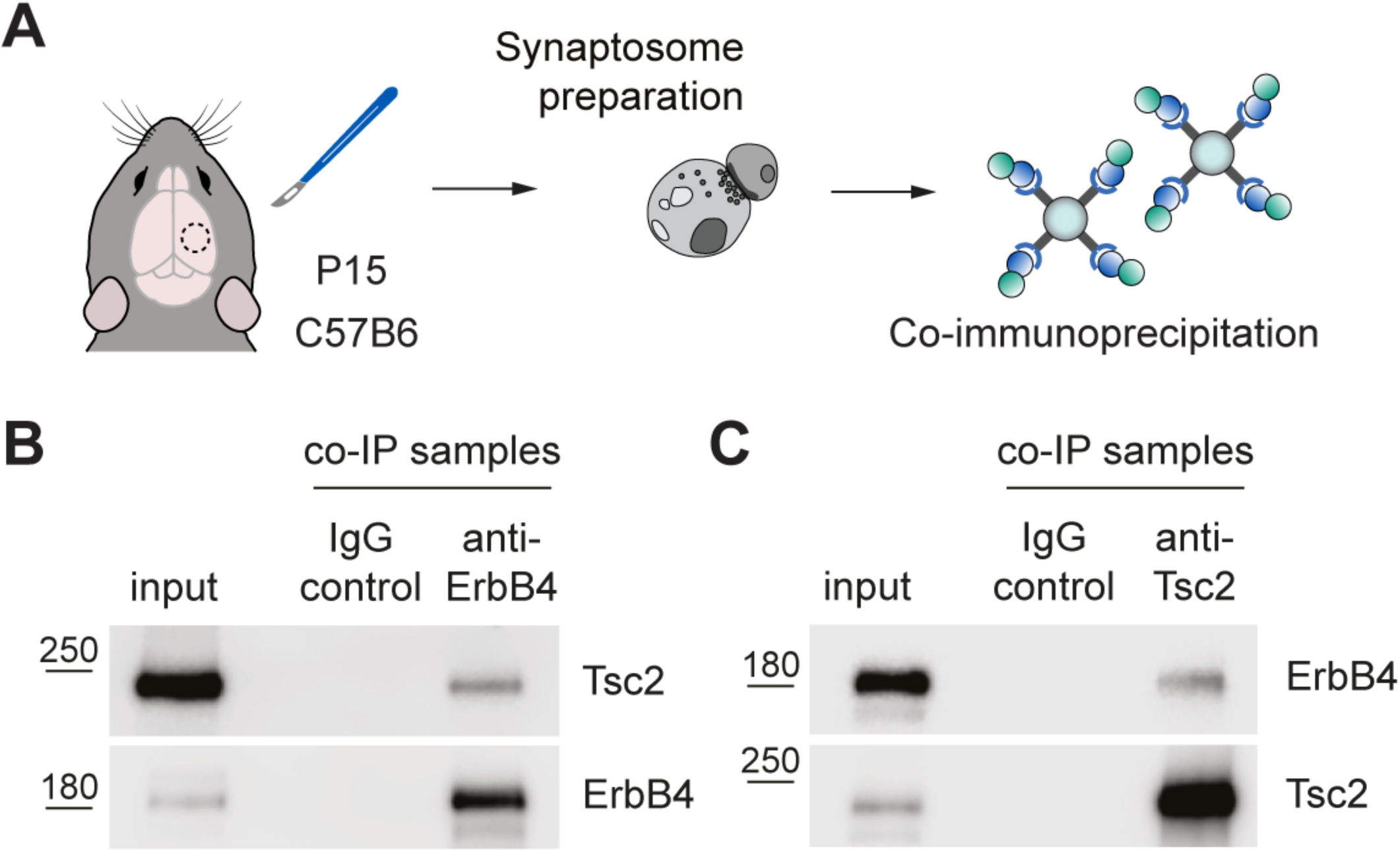
ErbB4 and Tsc2 interact at the synapse at the time of synaptogenesis. (**A**) Schematic of experimental design. (**B**) Western blot for Tsc2 and ErbB4 of anti-ErbB4 co-immunoprecipitation samples from cortical synaptosomes from P15 C57B6 mice (*n* = 3 independent co-immunoprecipitation experiments). (**C**) Western blot for Tsc2 and ErbB4 of anti-Tsc2 co-immunoprecipitation samples from cortical synaptosomes from P15 C57B6 mice (*n* = 3 independent co-immunoprecipitation experiments).

**Suppl. Fig. 5.**
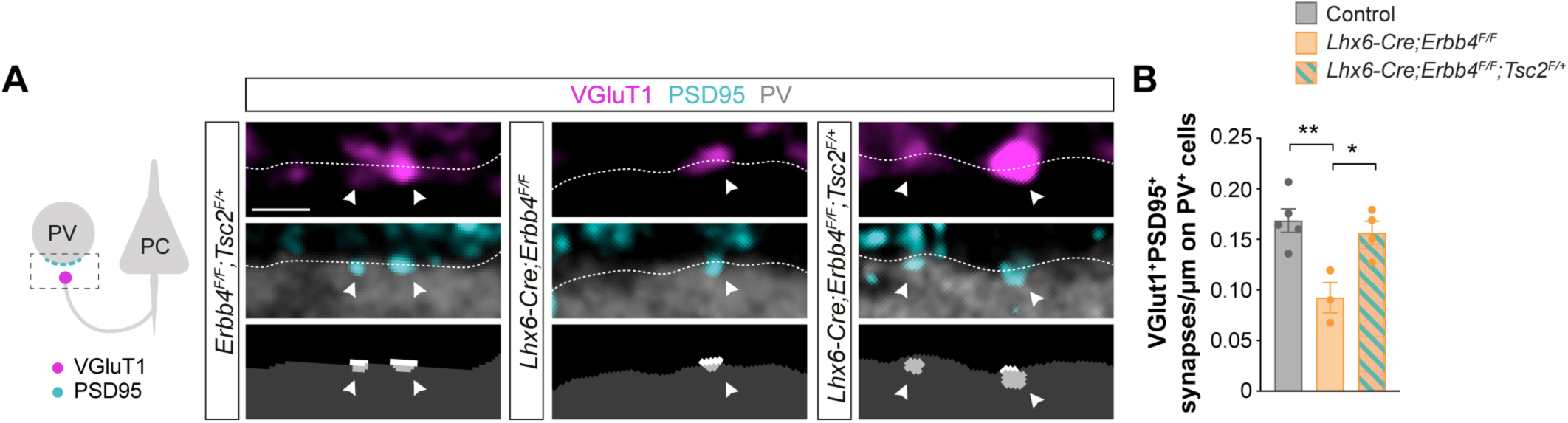
*Tsc2* deletion rescues synaptic loss in *Erbb4* conditional mutants. (**A**) Schematic of synaptic markers analyzed (left). Confocal images (top) and binary images (bottom) illustrating presynaptic VGluT1^+^ puncta (magenta) and postsynaptic PSD95^+^ clusters (cyan) in PV^+^ interneurons (grey) from P21 *Lhx6-Cre;Erbb4^F/F^;Tsc2-flox^F/+^* mice, *Lhx6-Cre;Erbb4^F/F^* and their control littermates. (**B**) Quantification of the density of VGluT1^+^PSD95^+^ synapses contacting PV^+^ interneurons (control, *n* = 124 cells from 5 mice; *Lhx6-Cre;Erbb4^F/F^*, *n* = 67 cells from 3 mice; *Lhx6-Cre;Erbb4^F/F^;Tsc2^F/+^*, *n* = 85 cells from 4 mice). ANOVA followed by Tukey’s multiple comparisons test: *P < 0.05, **P < 0.01. Data are mean ± s.e.m. Scale bar, 1 *µ*m.

**Suppl. Fig. 6.**
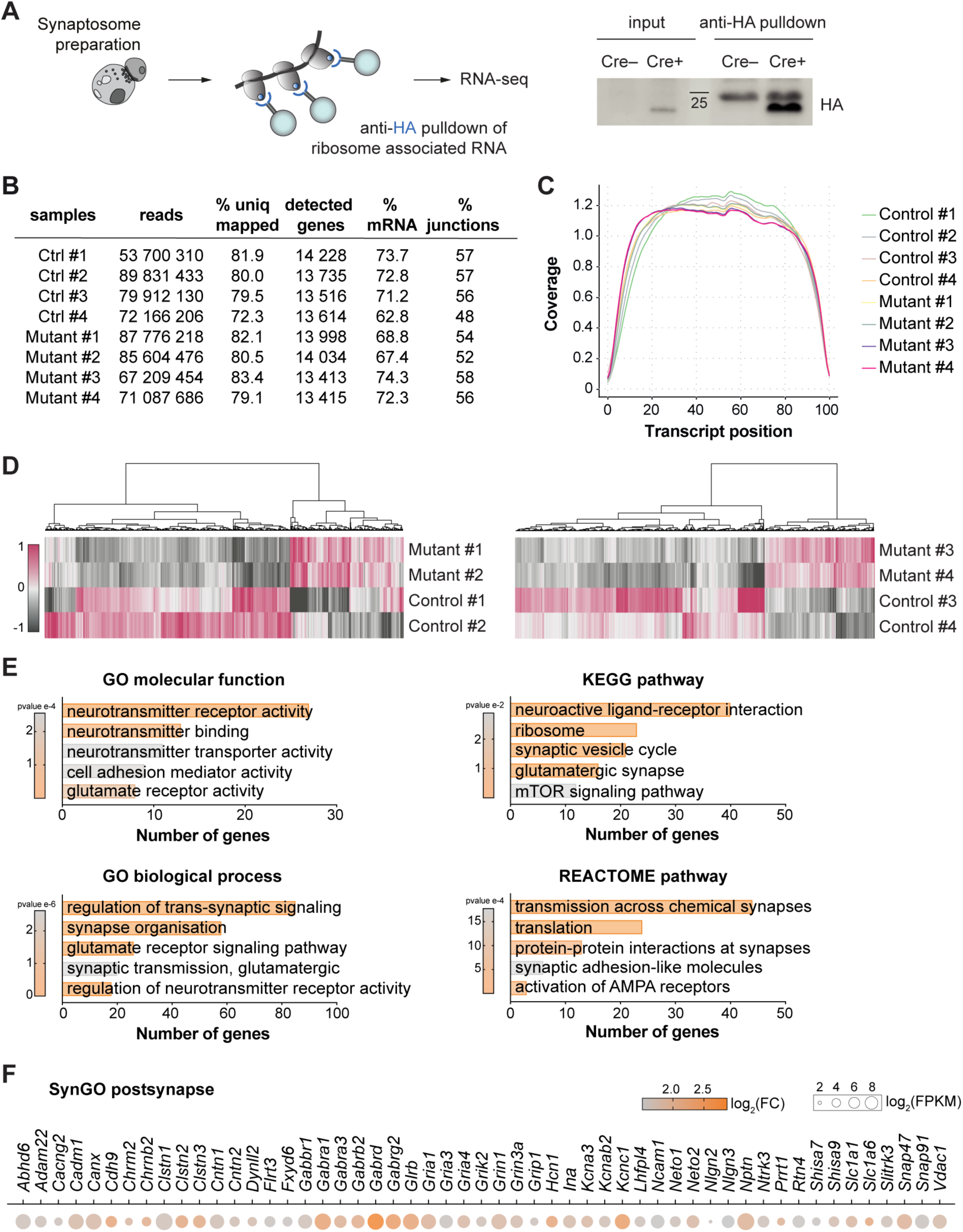
RNA sequencing data and gene ontology analysis of downregulated genes in *Erbb4* conditional mutants. (**A**) Schematic of experimental design (left). Western blot for HA of anti-HA pulldown samples from cortical synaptosomes from P15 *Lhx6-Cre;Rpl22^HA/HA^* mice (*n* = 2 independent co-immunoprecipitation experiments) (right). Note that the top band present in both Cre– and Cre+ anti-HA pulldown samples corresponds to IgGs from the pulldown antibodies. (**B**) FastQC analysis of RNA sequencing data for each replicate, showing total number of reads, % of reads uniquely mapped to the reference genome, number of detected genes, % of mRNA representation and % of reads mapped to exon-exon junctions. (**C**) 5’ to 3’ coverage plot showing the percentage of read bases across the transcript length. (**D**) Heatmaps showing significantly differentially expressed genes from two RNA sequencing batches from P15 *Lhx6-Cre;Erbb4^F/F^;Rpl22^HA/HA^* (conditional *Erbb4* mutants) cortical synaptosomes compared to *Lhx6-Cre;Erbb4^+/+^;Rpl22^HA/HA^* controls. Heatmap values are Deseq2 normalized counts. (**E**) Selected Gene Ontology (GO) terms, REACTOME and KEGG pathways significantly enriched in the dataset of downregulated genes in *Erbb4* mutants compared to controls. (**F**) Heatmap illustrating both control FPKM values and fold-change in conditional *Erbb4* mutants, for genes from “postsynaptic specialization” and “postsynaptic membrane” SynGO categories.

**Suppl. Fig. 7.**
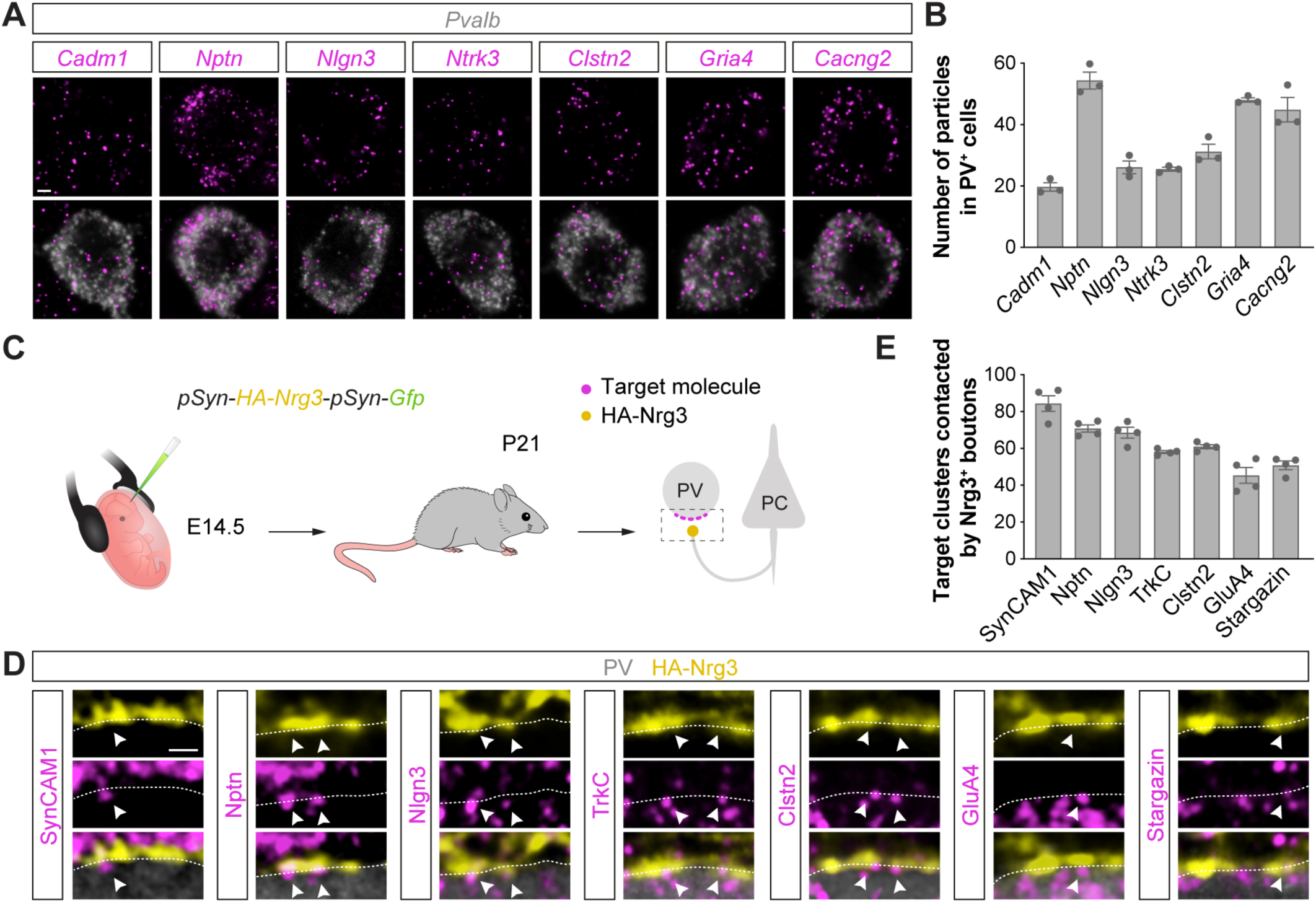
ErbB4 targets are expressed in developing cortical PV^+^ interneurons and are apposed to Nrg3^+^ excitatory synapses innervating PV^+^ cells. (**A**) Confocal images of target mRNAs (magenta) and *Pvalb* mRNA (grey) from single-molecule fluorescent in situ hybridization. (**B**) Number of mRNA particles per PV^+^ cell soma. *Cadm1*, *n* = 60 cells from 3 mice; *Nptn*, *n* = 66 cells from 3 mice; *Nlgn3*, *n* = 60 cells from 3 mice; *Ntrk3*, *n* = 60 cells from 3 mice; *Clstn2*, *n* = 60 cells from 3 mice; *Gria4*, *n* = 71 cells from 3 mice; *Cacng2*, *n* = 66 cells from 3 mice. (**C**) Schematic of experimental design. (**D**) Confocal images illustrating protein clusters (magenta) for the 7 targets at the surface of PV^+^ cell somas (grey), in close apposition to HA-Nrg3^+^ presynaptic boutons (yellow) from pyramidal cells electroporated with the plasmid *pSyn-HA-Nrg3-pSyn-Gfp* at embryonic day E14.5. (**E**) Proportion of target protein clusters contacted by Nrg3^+^ presynaptic boutons in PV^+^ interneurons. SynCAM1, *n* = 26 cells from 4 mice; Nptn, *n* = 26 cells from 4 mice; Nlgn3, *n* = 26 cells from 4 mice; TrkC, *n* = 26 cells from 4 mice; Clstn2, *n* = 26 cells from 4 mice; GluA4, *n* = 26 cells from 4 mice; Stargazin, *n* = 24 cells from 4 mice. Data are mean ± s.e.m. Scale bar, 3 *µ*m (A) and 1 *µ*m (D).

**Suppl. Fig. 8.**
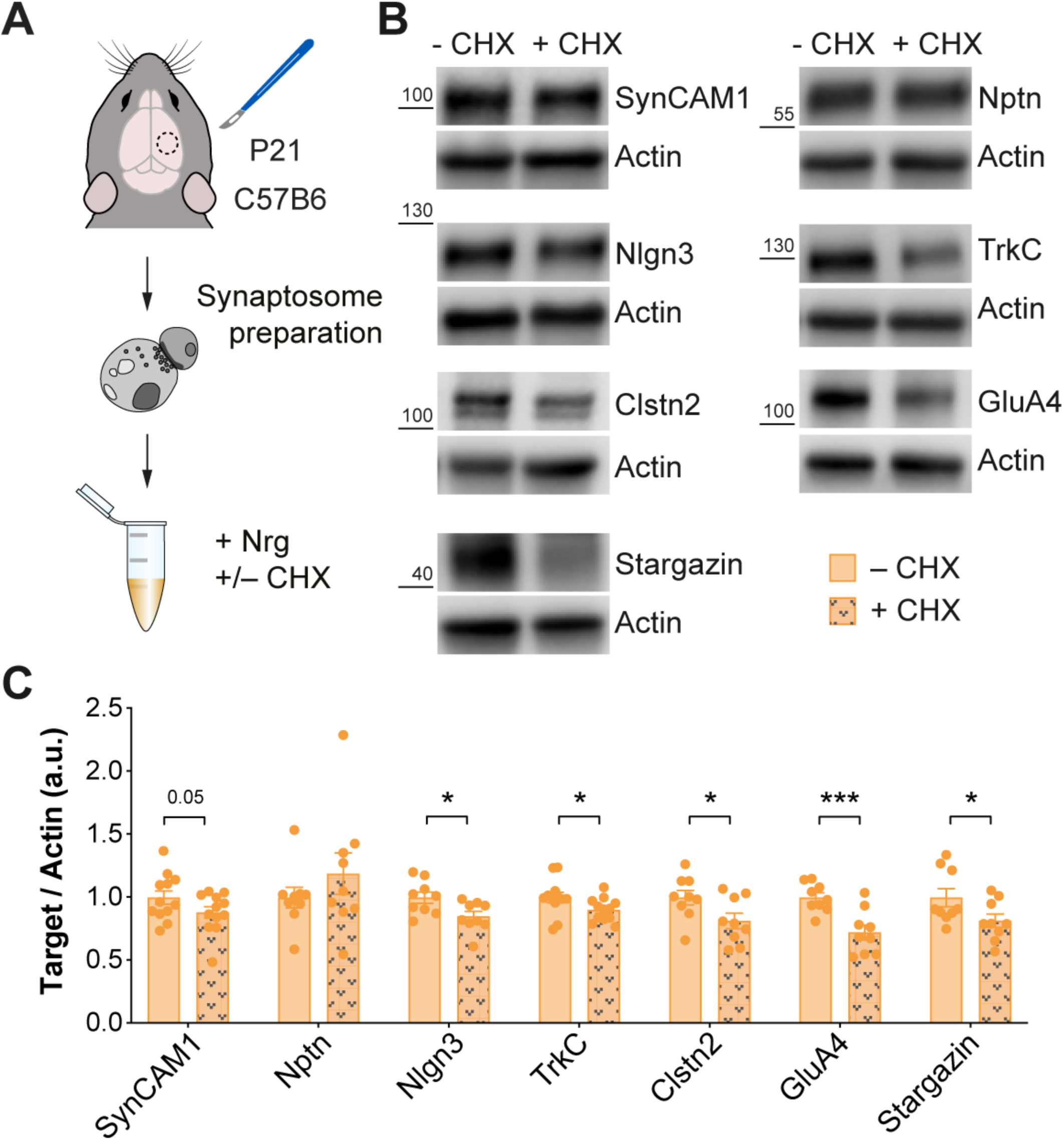
ErbB4 synaptic signaling regulates protein synthesis of synaptic proteins. (**A**) Schematic of experimental design. (**B**) Protein expression of SynCAM, Nptn, Nlgn3, TrkC, Clstn2, GluA4, Stargazin and actin assessed by Western blot of P21 cortical synaptic fractions treated with Neuregulin (Nrg) following pretreatment with or without cycloheximide (CHX). (**C**) Quantification of expression levels of SynCAM, Nptn, Nlgn3. TrkC, Clstn2, GluA4 and Stargazin normalized to actin. One-tailed Student’s unpaired t-tests: *P < 0.05, ***P < 0.001 (SynCAM1 and TrkC: –CHX *n* = 12 synaptosomes, +CHX, *n* = 12 synaptosomes; Nptn, Nlgn3, Clstn2, GluA4 and Stargazin: –CHX *n* = 9 synaptosomes, +CHX, *n* = 9). Data are mean ± s.e.m.

**Suppl. Fig. 9.**
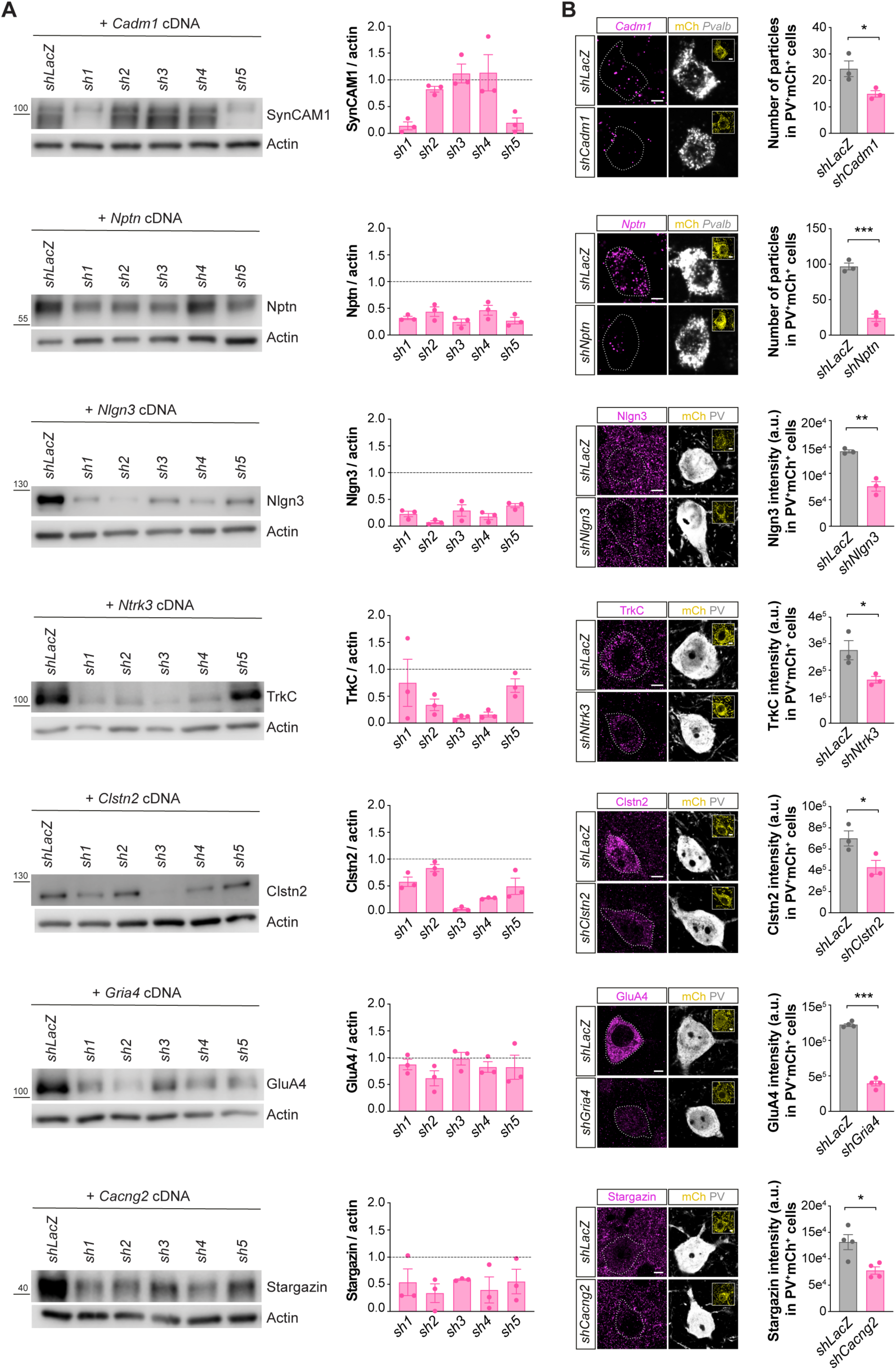
In vitro testing and in vivo validation of *shRNA* efficiency for downregulation of target targets. (**A**) In vitro protein expression assessed by Western blot of HA-tagged constructs of the 7 target targets from cells transfected with expression plasmids encoding the different targets and plasmids expressing *shRNAs* targeting the corresponding genes (left). Quantification of protein signal normalized to actin for each *shRNA*, relative to control transfections with a plasmid expressing a *LacZ*-targeting *shRNA* (*n* = 3 wells for each *shRNA*) (right). (**B**) Confocal images illustrating the knockdown of target RNAs (for *Cadm1* and *Nptn*) or target proteins (for Nlgn3, TrkC, Clstn2, GluA4 and Stargazin) in mCh^+^ (yellow) PV^+^ (grey) cells from P21 *Lhx6-Cre* mice injected with viruses expressing *shRNAs* targeting the genes of interest or with a control virus (*shLacZ*) (left). Quantification of RNA particles (for *Cadm1* and *Nptn*) or target protein staining intensity (for Nlgn3, TrkC, Clstn2, GluA4 and Stargazin) in mCh^+^ PV^+^ interneurons in knockdown and control mice (right). Two-tailed Student’s unpaired t-tests: *P < 0.05, **P < 0.01, ***P < 0.001 (*shCadm1*: *n* = 38 cells from 3 mice, *shLacZ*: *n* = 41 cells from 3 mice; *shNptn*: *n* = 46 cells from 3 mice, *shLacZ*: *n* = 55 cells from 3 mice; *shNgln3*: *n* = 59 cells from 3 mice, *shLacZ*: *n* = 44 cells from 3 mice; *shNtrk3*: *n* = 52 cells from 3 mice, *shLacZ*: *n* = 62 cells from 3 mice; *shClstn2*: *n* = 58 cells from 3 mice, *shLacZ*: *n* = 70 cells from 3 mice; *shGluA4*: *n* = 74 cells from 4 mice, *shLacZ*: *n* = 74 cells from 4 mice; *shCacng2*: *n* = 102 cells from 4 mice, *shLacZ*: *n* = 77 cells from 4 mice). Data are mean ± s.e.m. Scale bar, 5 *µ*m.

**Suppl. Fig. 10.**
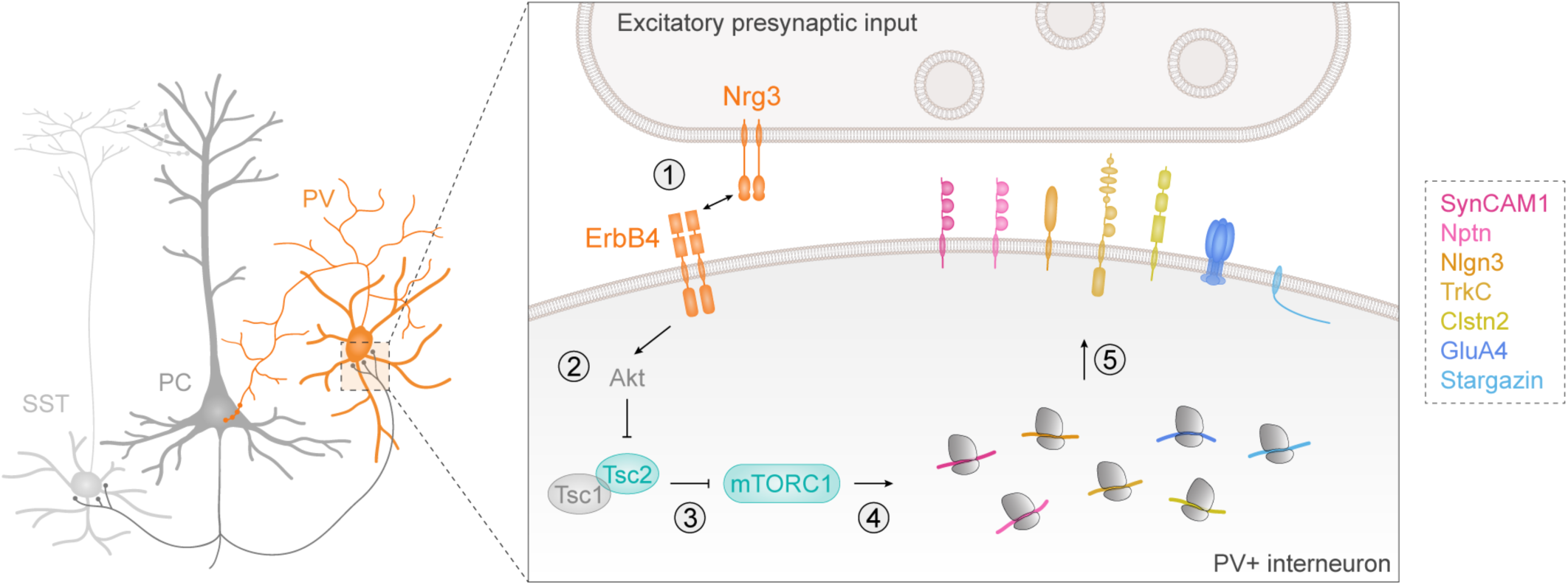
Summary of proposed signaling in developing cortical PV^+^ interneurons. Steps of ErbB4 synaptic signaling for the formation of excitatory synapses received by cortical PV^+^ interneurons: (1) activation of ErbB4 through binding to Nrg3 ligand, (2) inhibition of Tsc2 through Akt-dependent phosphorylation cascade, (3) inhibition of mTORC1 through Rheb-dependent phosphorylation cascade, (4) activation of synaptic protein synthesis of ErbB4 targets and (5) excitatory synapse formation through ErbB4 targets during postnatal development.

